# Biophysical properties of intermediate states of EMT outperform both epithelial and mesenchymal states

**DOI:** 10.1101/797654

**Authors:** Yoran Margaron, Tomoaki Nagai, Laurent Guyon, Laetitia Kurzawa, Anne-Pierre Morel, Alice Pinheiro, Laurent Blanchoin, Fabien Reyal, Alain Puisieux, Manuel Théry

**Affiliations:** Université de Paris, CEA/ INSERM/AP-HP, Institut de Recherche Saint Louis, UMR976, HIPI, CytoMorpho Lab, Hopital Saint Louis, 1 Avenue Claude Vellefaux, 75010 Paris, France; Université Grenoble-Alpes, CEA/INRA/CNRS, Interdisciplinary Research Institute of Grenoble, UMR5168, LPCV, CytoMorpho Lab, 17 rue des Martyrs, 38054 Grenoble, France; University Grenoble Alpes, CEA/INSERM, Interdisciplinary Research Institute of Grenoble, Biology of Cancer and Infection UMR_S 1036 38000 Grenoble, France; Université de Lyon, Université Claude Bernard Lyon 1, INSERM 1052, CNRS 5286, Centre Léon Bérard, Cancer Research Center of Lyon, Lyon, France; Residual Tumor & Response to Treatment Laboratory (RT2Lab), PSL Research University, Translational Research Department, F-75248, Paris, France

## Abstract

Potential metastatic cells can dissociate from a primary breast tumor by undergoing an epithelial-to-mesenchymal transmission (EMT). Recent work has revealed that cells in intermediate states of EMT acquire an augmented capacity for tumor-cell dissemination. These states have been characterized by molecular markers, but the structural features and the cellular mechanisms that underlie the acquisition of their invasive properties are still unknown. Using human mammary epithelial cells, we generated cells in intermediate states of EMT through the induction of a single EMT-inducing transcription factor, ZEB1, and cells in a mesenchymal state by stimulation with TGFβ. In stereotypic and spatially-defined culture conditions, the architecture, internal organization and mechanical properties of cells in the epithelial, intermediate and mesenchymal state were measured and compared. We found that the lack of intercellular cohesiveness in epithelial and mesenchymal cells can be detected early by microtubule destabilization and the repositioning of the centrosome from the cell-cell junction to the cell center. Consistent with their high migration velocities, cells in intermediate states produced low contractile forces compared with epithelial and mesenchymal cells. The high contractile forces in mesenchymal cells powered a retrograde flow pushing the nucleus away from cell adhesion to the extracellular matrix. Therefore, cells in intermediate state had structural and mechanical properties that were distinct but not necessarily intermediate between epithelial and mesenchymal cells. Based on these observations, we found that a panel of triple-negative breast cancer lines had intermediate rather than mesenchymal characteristics suggesting that the structural and mechanical properties of the intermediate state are important for understanding tumor-cell dissemination.

## Introduction

In multicellular organisms, interconversion between epithelial and mesenchymal phenotypes through the process of epithelial-to-mesenchymal transition (EMT) provides the cell plasticity required during critical steps of embryogenesis (Thiery et al., 2009). This reversible program involves profound changes in cell morphology and behavior, enabling cells to migrate to distant sites in the embryo and participate in the formation of internal organs. In pathological conditions, EMT has been implicated in tissue fibrosis as well as in several steps of tumor development and progression, including tumor initiation, invasion, metastatic dissemination and resistance to therapy (Dongre and Weinberg, 2019). Recent lines of evidence suggest that these distinct steps are not supported by a unique cell state. In contrast to the classic view of EMT as a binary process with two exclusive phenotypes, either fully epithelial of fully mesenchymal, the program entails epithelial cells entering into a variety of intermediate states with different functions and properties (Gupta et al., 2019; Nieto et al., 2016; Pastushenko and Blanpain, 2019). Consistent with this notion, cancer cells exhibiting a hybrid epithelial-mesenchymal phenotype have been identified in both primary tumors and at sites of dissemination (Aceto et al., 2014; Baccelli et al., 2013; Kröger et al., 2019; Yu et al., 2013). In skin and mammary primary tumors, cells in intermediate states of EMT have been shown to localize at the invasive front (Pastushenko et al., 2018). Overall, these data suggest that EMT-associated pliancy is a prominent source of intra-tumor phenotypic and functional heterogeneity (Puisieux et al., 2018). Although transcriptional and epigenetic landscapes of different EMT intermediate states have been characterized recently (Pastushenko et al., 2018), we still lack a coherent overview of the intracellular organization and mechanical properties of these states.

The shape, size, and position of organelles are key regulators of cell functions (Bornens, 2008). The cytoskeleton controls cell shape and organizes the entire intracellular space, from the cytoplasm to the architecture of the nucleus (Blanchoin et al., 2014; Lele et al., 2018; Mimori-Kiyosue, 2011; Uhler and Shivashankar, 2017). The microtubule network organizes the endo-membrane network and direct intracellular transport (de Forges et al., 2012). In turn the microtubule network is organized by the centrosome, the position of which is the outcome of a broad integration process that balances mechanical forces and responds to numerous signaling pathways (Bornens, 2008; Nigg, 2014). Indeed, the position of the centrosome rapidly responds to fine changes in cell adhesion or the activation of surface receptors. By doing so, centrosome repositioning powers massive structural changes including cell polarization and cilium growth, endo- and exocytosis and lumen formation, and thus constitutes a key readout to interpret the functional state of a cell (Bornens, 2012; Stinchcombe and Griffiths, 2014; Tang and Marshall, 2012). In parallel, the actin network powers changes in cell shape and intracellular architecture. In particular, the actin network is directly connected to the nucleus, and can act as a structural hub that transmits mechanical constraints on the nucleus (Alam et al., 2015; Gomes et al., 2005; Luxton et al., 2010). Hence mechanical forces not only position the nucleus but also affect chromatin organization and gene expression, notably during cell differentiation (Uhler and Shivashankar, 2017). Therefore, the position of the nucleus can be considered as another key readout of the mechanical and differentiation state of a cell. Given that the progression through multiple stages of EMT involves numerous changes in cell adhesion, reprogramming of gene expression and the acquisition of particular migration properties, we reasoned that organelle positioning and intracellular mechanical forces could reveal how these changes are associated with, and possibly powered by, specific reorganizations of intracellular architectures.

In vitro 3D modelling of complex physiological behaviors is now becoming technological feasible for processses such as symmetry breaking, lumen formation, the patterning of cell differentiation and the formation of invasive tissue outgrowth (Vianello and Lutolf, 2019; Wan, 2016). However, the final shape of these large multicellular arrangements is the outcome of numerous processes that involve the regulations of intercellular interactions, cell shape and migration, the respective contributions of which are difficult to entangled. In addition, the 3D conformation is not optimal for intracellular imaging. Therefore to identify the key elementary processes regulating cell behaviors, a trade-off has to be made between tissue-like morphological mimicry and focused biophysical investigation. Micro-engineering of cell-culture devices now offer the possibility to impose defined spatial boundary conditions at the multi-cellular, cellular or sub-cellular level, which can recapitulate specific geometrical and physical constraints cells are submitted to in vivo (Laurent et al., 2017; Théry, 2010). Thereby it is possible to induce specific morphogenetic events in controlled and reproducible conditions. In addition, these approaches have offered the possibility to perform automated, rapid and precise image acquisition and analysis. Here we used 2D surface micropatterning to direct the self-organization of human mammary epithelial cells (HMEC) in simple and stable conformations in order to study in detail, the changes in the internal architecture of the cell in an epithelial or mesenchymal state or in an intermediate epithelial/mesenchymal state.

## Results

### Establishing a model for intermediate stages of EMT

To investigate the variations in the positioning of intracellular organelles at various stages of EMT, we used normal human mammary epithelial cells (HMECs) because these cells are non-transformed and present most features of primary epithelial cells including a limited lifespan (Lindley and Briegel, 2010). HMEC can adopt an advanced mesenchymal state and express a panel of EMT-associated transcription factors after exposure to 5 ng/mL of TGFβ for 5 days (Lindley and Briegel, 2010). A transcriptomic analysis revealed that ZEB1 is the first EMT transcription factor to be upregulated upon exposure to TGFβ; and its RNA levels increase by 10 fold within 12 hours (Lindley and Briegel, 2010). Interestingly, the expression of only ZEB1 is necessary and sufficient to confer cells with the characteristics of intermediate stages of EMT (Krebs et al., 2017; Liu et al., 2014; Morel et al., 2012; Stemmler et al., 2019). Therefore, to specifically investigate the intermediate state of EMT, we engineered a Tet-On–inducible HMEC line to over-express ZEB1 (HMEC-ZEB1) through 24-hour exposure to doxycycline in the culture medium. Hence the intermediate and complete states of EMT were compared by evaluating doxycycline-stimulated HMEC-ZEB1s with TGFβ–stimulated MCF10A cells.

After 24-hour exposure to doxycycline, HMEC-ZEB1s lost their capacity to grow in multicellular islands (Figure 1a). However, the presence of some cell aggregates suggested that the cells maintained a certain level of cohesiveness, which was preserved even after 5 days of doxycycline exposure. By contrast, cell exposure to 5 ng/mL of TGFβ led to a complete scattering of cells after 5 days (Figure 1a). We further characterized the states of these cells by measuring EMT markers. At the mRNA level, we confirmed that a 24-hour treatment with doxycyclin induced a strong over-expression of ZEB1, but found no noticeable effect on ZEB2 or SNAI1 (Supplementary Figure 1a). The amount of ZEB2 transcripts increased after 70 hours but remained an order of magnitude below the amount of ZEB1. The amount of protein followed the same trend: ZEB1 appeared overexpressed after 24-hour exposure to doxycyclin, whereas ZEB2 and SNAIL were unaffected. As expected, the concentrations of the three transcription factors were highly increased by a 5-day exposure to TGFβ (Figure 1b). Variations of the amounts of E-cadherin and vimentin were consistent with this trend: 5-day exposure to TGFβ, the expression level of E-cadherin was strongly diminished and vimentin increased, whereas over-expression of ZEB1 for 24 hours or 5 days induced only milder changes (Figure 1c). We concluded that HMEC over-expressing ZEB1 were maintained in a steady and intermediate state of EMT. These results were used to define three states of EMT as follows: non-treated HMEC cells represented the epithelial state (i.e E cells); HMEC expressing ZEB1 for 24 hours represented a stable intermediate state (i.e E/M cells); and HMEC cells treated with TGFβ for 5 days represented the mesenchymal state (i.e. M cells).

**Figure 1:**
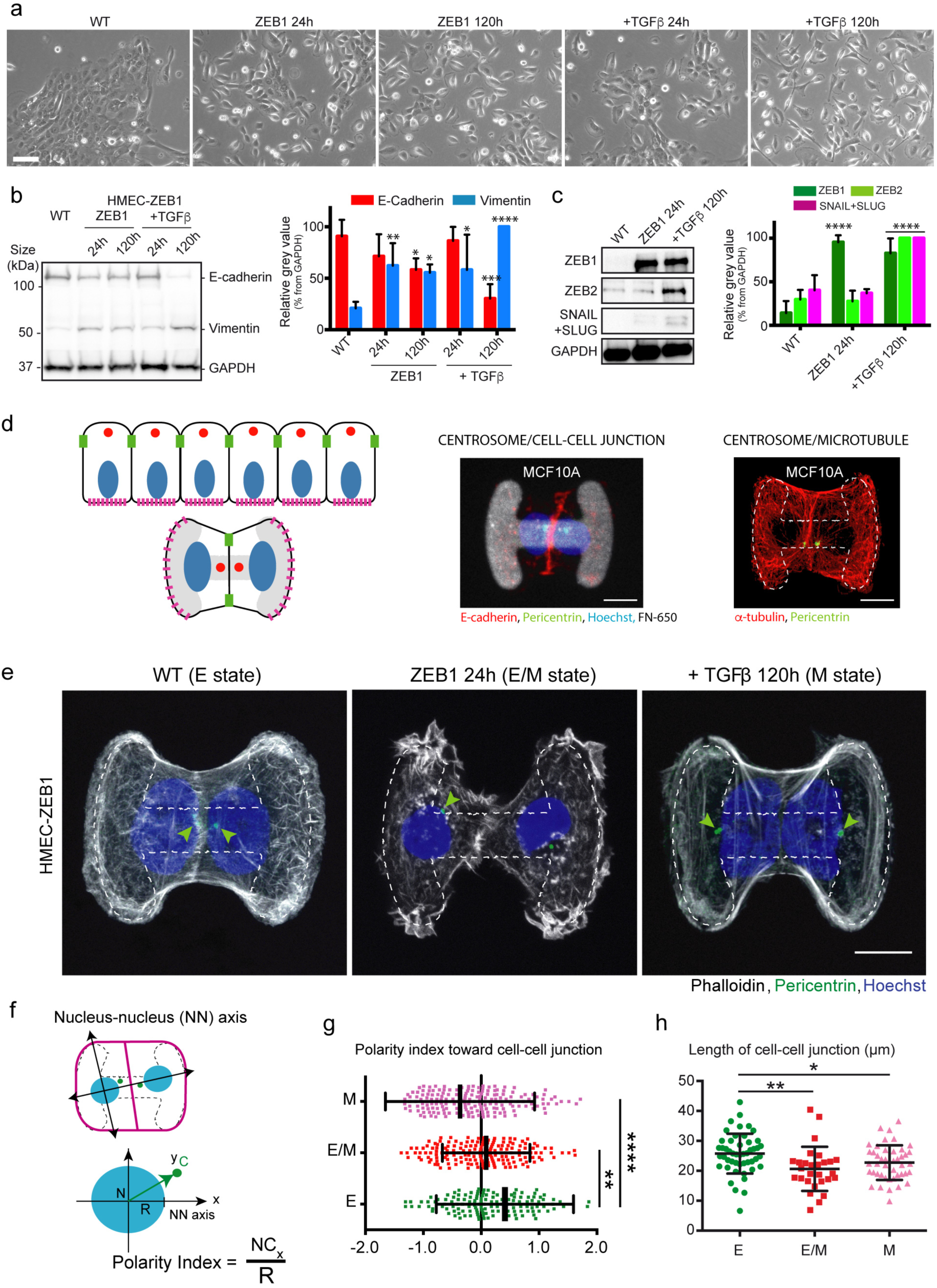
Polarity differences of cells in the epithelial (E) state, intermediate (E/M) state and mesenchymal (M) state. **a** Representative images of HMEC-ZEB1s cultured alone (WT; E state; left panel), with exposure to doxycycline (ZEB1 overexpression; E/M state; middle panel) for 24 hours or 5 days, or with exposure to TGFβ (M state; right panel) for 24 hours or 5 days. Scale bars represent 30 µm. **b** Expression of ZEB1, ZEB2, SNAIL/SLUG in HMEC-ZEB1s cultured alone (WT), with exposure to doxycycline (ZEB1 overexpression) for 24 hours, or with exposure to TGFβ for 5 days. Quantification of ZEB1, ZEB2 and SNAIL/SLUG protein expression, relative to GAPDH, in WT, doxycycline- and TGFβ-treated cells. ****: p<0.0001 by two-way ANOVA and Fisher’s LSD multiple comparison test. Error bars indicate SD, N=3 independent experiments. **c** Expression of vimentin and E-cadherin in HMEC-ZEB1s cultured alone (WT), with with exposure to doxycycline (ZEB1 overexpression) for 24 hours or 5 days, or with exposure to TGFβ for 24 hours or 5 days. Right panel shows a quantification of vimentin and E-cadherin protein expression, relative to GAPDH, in WT, ZEB1, and TGFβ HMEC-ZEB1s. *: p<0.05, **: p<0.01, ***: p<0.001, ****: p<0.0001 by two-way ANOVA and Fisher’s LSD multiple comparison test. Error bars indicate SD, N=3 independent experiments. **d** Schematic representation of epithelial-cell doublets cultured on H shape micropattern (in grey) recapitulating the intracellular organization of cells in a tissue (left panel). Images of the intracellular organization of MCF10A cells cultured on H micropatterns (middle and right panels). Middle panel shows E-cadherin (red), pericentrin (green) and Hoechst (blue) for cell-cell junction, centrosome and nuclei location respectively. Fluorescent fibronectin coating of the micropattern is displayed in grey. Scale bars represent 20 μm. **e** Representative images showing HMEC-ZEB1 doublets on H-shaped micropatterns (dashed lines). HMEC-ZEB1s cultured alone (WT), with exposure to doxycycline (ZEB1 overexpression) for 24 hours, or with exposure to TGFβ for 5 days.. Cells were stained for F-actin (white), pericentrin (green) and Hoechst (blue). Arrowheads highlight centrosome positioning. Scale bar represents 20 µm. **f** Measurements of centrosome position. The scheme shows the centrosome coordinates with respect to the center of mass of the nucleus: x-axis corresponds to the nucleus-nucleus (NN) axis passing through the center of two nuclei. The distance from nucleus center to centrosome is normalized by nucleus radius. **g** Quantification of the polarity indeces for cells cultured under the conditions described in (c) in E, E/M and M cells. **: p<0.01, ****: p<0.0001, by Student *t*-test. Error bars indicate SD, n>296 cells, N=3 independent experiments. **h** Quantification of the length of the cell-cell junction in E, E/M and M cells. *: p<0.05, **: p<0.01, by Student *t*-test. Error bars indicate SD, n>56 cells, N=3 independent experiments.

### Specific remodelling of intracellular organisation in partial and complete EMT

To identify whether cells in various states of EMT adopt distinct intracellular architectures, HMEC in E-, E/M- and M-states were cultured as cell doublets on H-shaped fibronectin-coated micropatterns over 24 hours. The geometry of the micropattern dictated cell shape and positioning, with the intercellular junction situated at the vertical axis bisecting the H and the cell-matrix adhesions distal at the respective vertical bars of the H (Tseng et al., 2012). The coexistence but segregation of cell-cell adhesions and cell-ECM adhesions captured the minimal set of conditions to recapitulate the epithelial configuration in 2D model (Burute and Thery, 2012)(Figure 1d). Furthermore, by imposing reproducible and comparable cell conformations, these conditions permit systematic quantifications of organelle positioning (Schauer et al., 2010; Théry et al., 2006) including the repositioning of the centrosome and nucleus with TGFβ-induced EMT in MCF10A cell line (Burute et al., 2017).

Non-stimulated HMEC-ZEB1s and the parental HMECs, both in the E state, adopted the same shape and position on H-shaped micropatterns as previously observed in MCF10A (Burute et al., 2017) (Figure 1e, Supplementary Figure S2a). The relative position of centrosome with respect to the center of mass of the nucleus and the position of the intercellular junction was captured as an internal polarity index of the cell (Figure 1f). Cells in the E state displayed the characteristic epithelial configuration, with a positive polarity index characteristic of the nucleus-centrosome axis being oriented toward the intercellular junction (Burute et al., 2017)(Figure 1e-left and Figure 1g). Cells in the M-state, treated for 5 days with TGFβ prior to plating on the micropatterns, displayed the expected complete reversal of the nucleus-centrosome axis, being oriented towards the cell-matrix adhesion and with a negative polarity index (Figure 1e-right and Figure 1g). Interestingly, the architecture of cells in the E/M-state was distinct from the E-state and M-state: as with the M-state, the intercellular junction was shorter than in the E-state (Figure 1e and 1h), but unlike the M-state, the nucleus-centrosome axes were randomly oriented, as revealed by the polarity index being near zero (Figure 1g). By way of controls, for the parental HMEC line, doxycycline had no effect on cell shape and polarity index, and 5-day treatment with TGFβ induced nucleus-centrosome axis reversal (Supplementary Figure S2b).

To reveal the specific repositioning of each organelle in E-, E/M- and M-state cells, we plotted the spatial distribution of nuclei and centrosomes in those three conditions (Figure 2a). The position of these organelles were visualized and quantified using probability density maps (Schauer et al., 2010)(Figure 2b). For the comparison of single pairs of probabilistic density maps, we used the two-sample kernel density-based test (Duong et al., 2012). Organelle positions were also quantified by measuring their distance to the cell center of mass (Figure 2c). These analyses showed that centrosomes repositioned from the intercellular junction to the cell center as cells transit from E to E/M, and that nuclei repositioned from cell center to intercellular junction as cells transit from E/M to M state.

**Figure 2:**
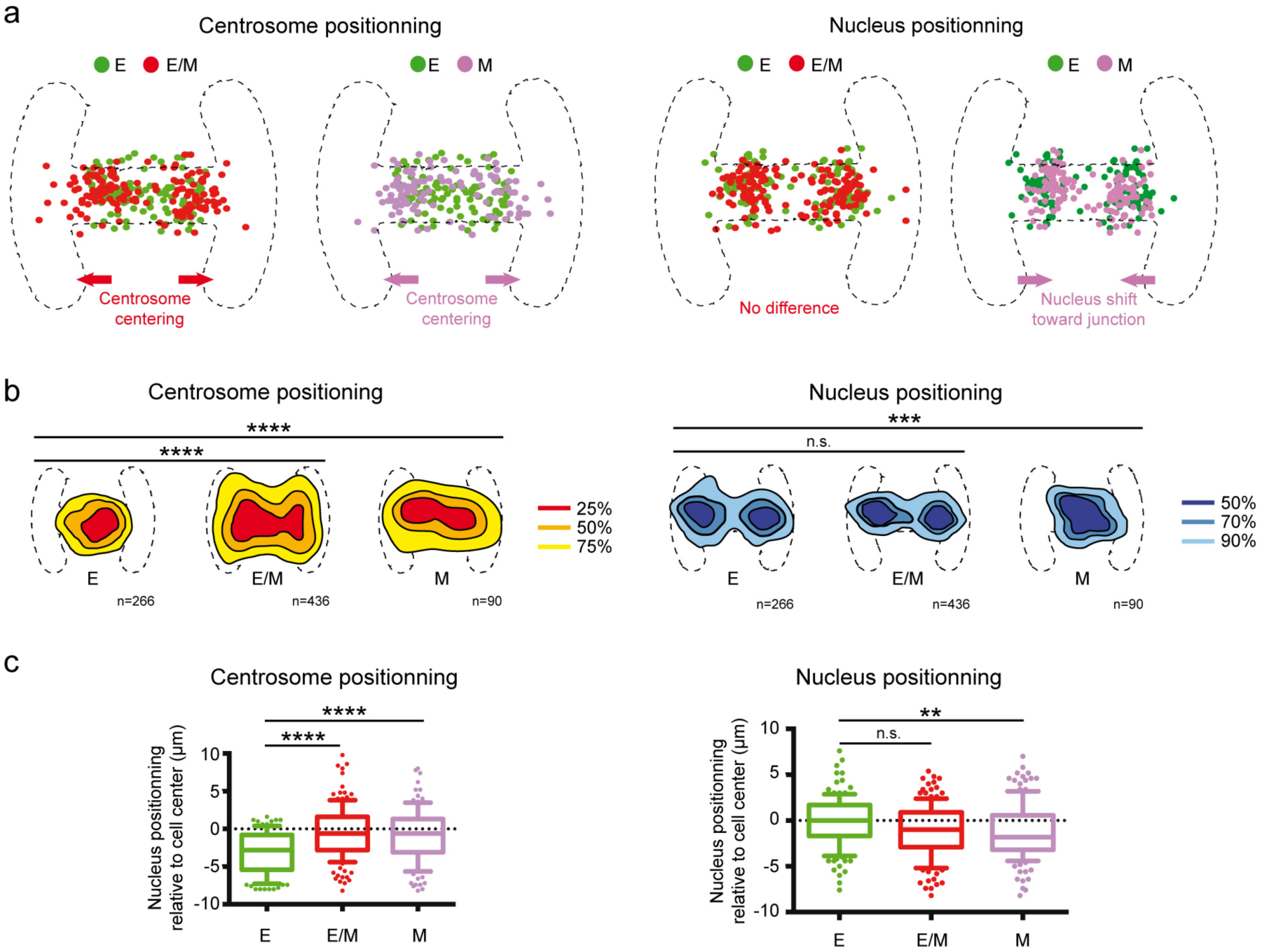
Nucleus and centrosome positioning in E, E/M and M cells. a Graphs showing the distributions of centrosomes (left two panels) and nuclei (right two panels) in HMEC doublets cultured on H micropatterns. Distributions in E/M and M cells were overlaid with distribution in E cells to facilitate visual comparison. b Probabilistic density maps representing the regions in which centrosomes (left panels in red/yellow) and nuclei (right panel in dense/light blue) are found in cell doublets from the analysis of a minimum of 90 cells per condition. ***: p<0.001, ****: p<0.0001 from two-sample kernel density-based test. N=3 independent experiments. c Quantification of the distance between centrosomes (left panel) or nuclei (right panel) and center of mass of the cell in E, E/M and M cells. **: p<0.01, ****: p<0.0001 by Kruskal-Wallis test and Dunn’s multiple comparison test. n>90 cells, N=3 independent experiments.

### Microtubules destabilisation and centrosome centring in E/M and M cells

We then investigated the molecular and cellular mechanisms regulating the sequential repositioning of the centrosome and the nucleus as cells progress through EMT. Microtubule stability was evaluated in E, E/M and M cells. given that centrosome positioning strongly depends on microtubule dynamics (Elric and Etienne-Manneville, 2014; Letort et al., 2016; Pitaval et al., 2017). Our previous modeling and experimental data showed that dynamic microtubules apply balanced forces on cell edges and maintain the centrosome at the cell center, whereas microtubule stabilization tends to break network symmetry and push the centrosome away from the cell center (Burute et al., 2017). Therefore, cells were briefly exposed to a cold treatment (15 minutes, 4°C) to depolymerize dynamic but not stable microtubules (Jones et al., 1980). In E cells, large proportion of microtubules resisted exposure to 4°C. By contrast, in E/M or M cells, only a very few microtubules resisted exposure to 4°C (Figure 3a and b). By way of controls, for the parental HMEC line, doxycycline had no effect and the 5-day treatment with TGFβ also destabilized microtubules (Supplementary Figure S2c). These results showed that microtubules were less stable in E/M or M cells than in E cells, which was consistent with centrosome being more central in E/M and M cells and off-centered in E cells.

**Figure 3:**
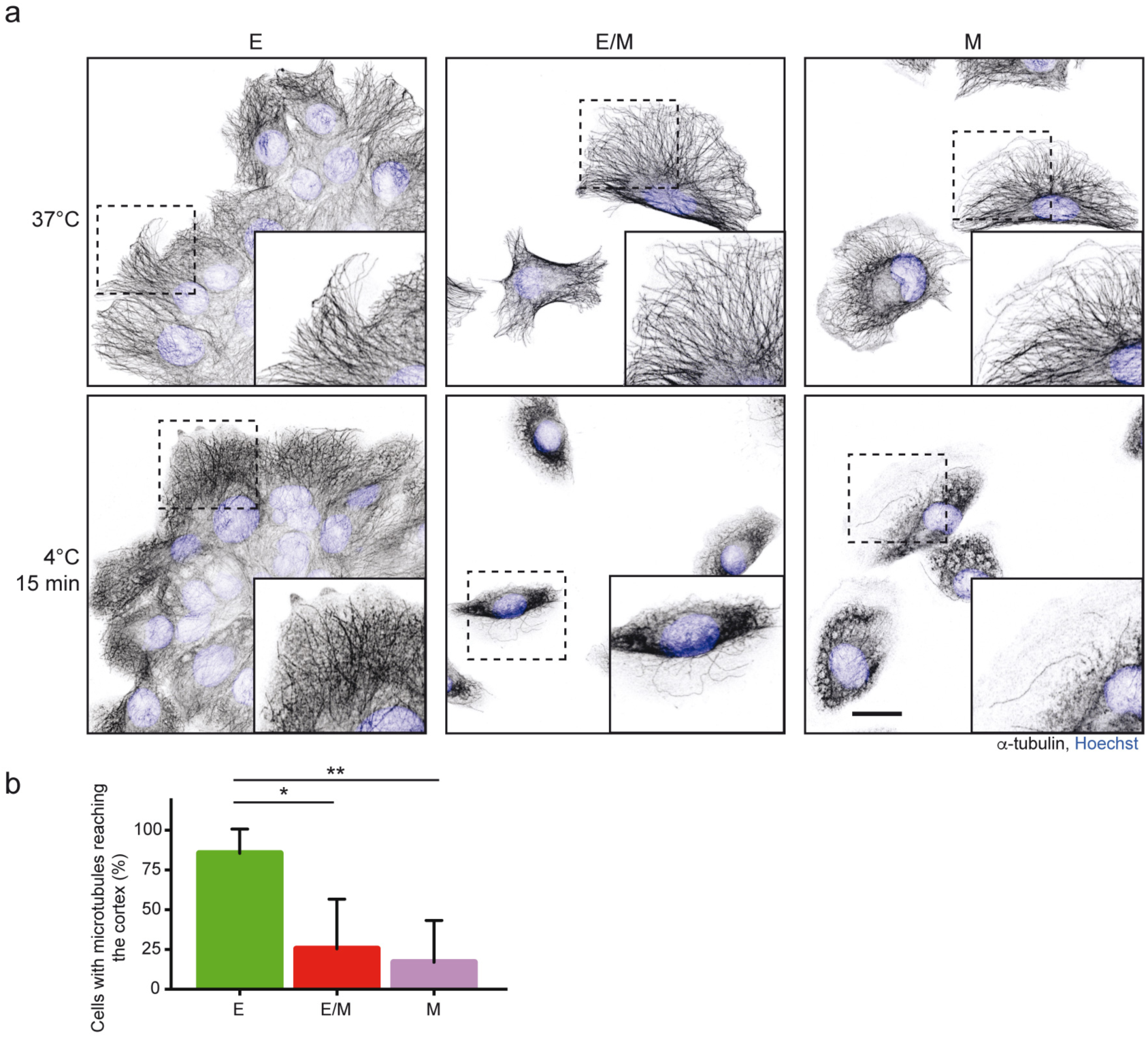
EMT induced microtubule destabilization. a Images show microtubule networks in cells in the epithelial (E) state, epithelial/mesenchyme (E/M) state and mesenchymal (M) state in normal culture conditions (top) or after a 15-minute exposure to cold treatment (4°C) in order to reveal the more stable microtubules. The cells were stained for a The cells were stained for table microtubuleswere n ZEB1 induced E/M and TGFght panel) and cell center of mass in itµm. b Quantification of the number of cells in which microtubules were identified at the cell cortex after cold treatment. *: p<0.05, **: p<0.01 by one-way ANOVA and Tukey’s multiple comparison test. Error bars indicate SD, n>50, N=3 independent experiments.

### Acto-myosin activity regulates nucleus decentering in M cells

Actin network dynamics were evaluated in E, E/M and M cells, because the nucleus is tightly associated with actin filaments (Lele et al., 2018; Luxton et al., 2010) and in mesenchymal-like migration, the nucleus is positioned away from focal adhesions and near the rear of the cell due the retrograde flow of contractile actin bundles generated along focal adhesions (Dupin et al., 2009; Gomes et al., 2005). Actin filaments were imaged in real time in live cells over a 30-minute period using SiR-actin (Figure 4a). Measurements of the displacement of actin bundles revealed that in M cells, the actin flow at the height (z-plane) of the nucleus was 3-fold greater than that in E and E/M cells (Figure 4b). Given that the retrograde flow of actin bundles is generally powered by acto-myosin contraction, and that actin bundles appeared thicker and straighter in M cells (Figure 4a), we decided to then investigate myosin activity and its contribution to organelle positioning.

**Figure 4:**
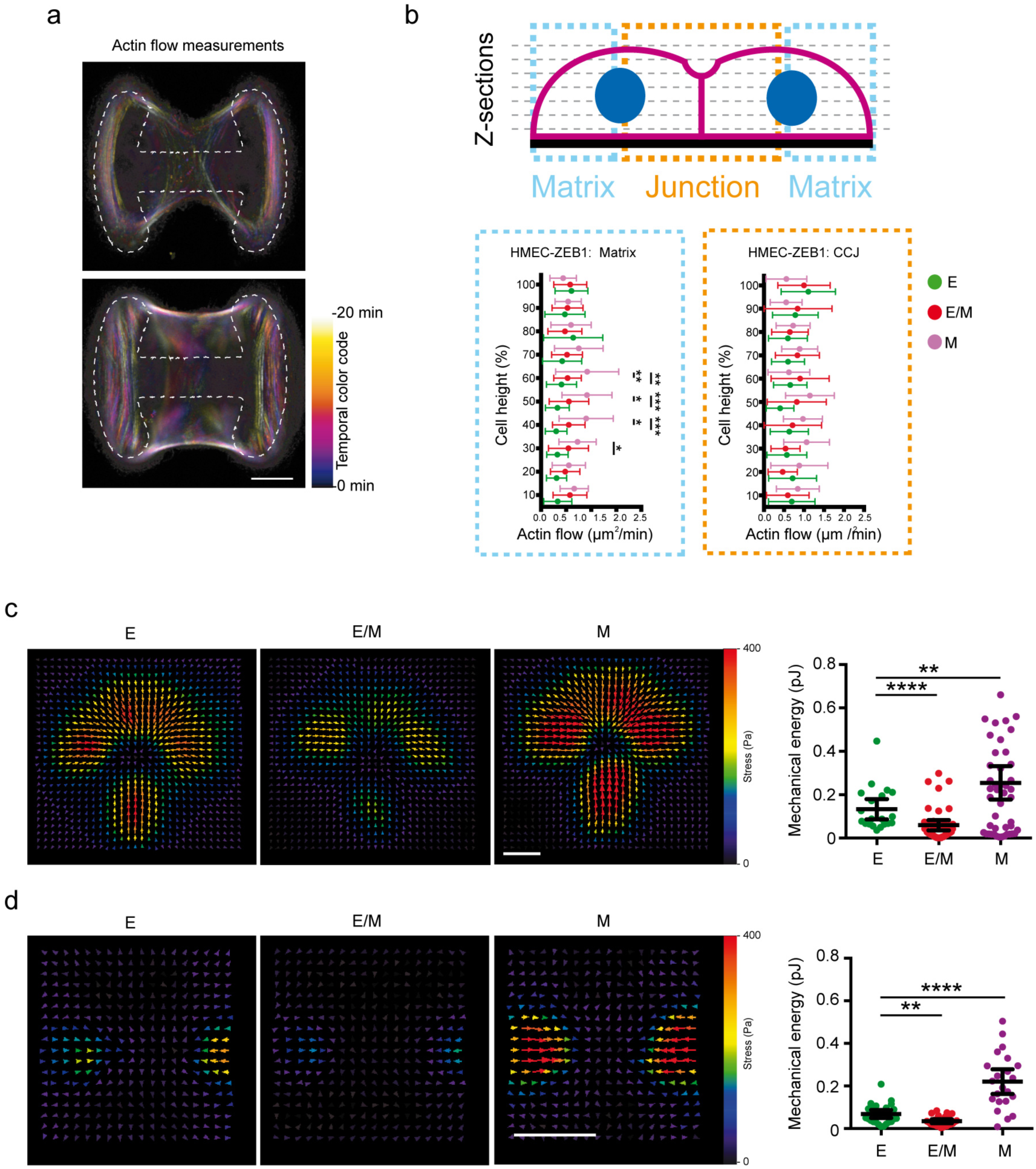
Actin retrograde flow and cell traction forces in E, E/M and M cells. a Overlayed time-sequence images (−20 to 0 minutes) in distinct colors to illustrate actin flow in E/M and M cells. b Quantification of actin flow at various cell heights in cells stained with SiR-actin and cultured on H-shape micropatterns. *: p<0.05, **: p<0.01, ***: p<0.001 from Mann-Whitney test. n>12, error bars indicate SD, N=3 independent experiments. c-d. Averaged traction-stress field (in Pascal) in E, E/M and M cells cultured for 24 hours on crossbow (c) and rectangle (d) micropatterned polyacrylamide hydrogels. n=20, N=2 independent experiments. The graph displays the mechanical energy of cell traction in each condition. n>20, error bars indicate 95% confidence interval, N=2 independent experiments. **: p<0.01, ****: p<0.0001 by Mann Whitney U test. Scale bar represents 20µm.

First, we evaluated the tensional state of HMECs by measuring the traction forces produced by individual cells on a fibronectin-coated micropattern. A crossbow micropattern was chosen for a single cell, to mimic the geometrical constraints imposed on one of the two individual cells in the H-micropattern. The shape of a crossbow imposed a continuous adhesive edge and two non-adhesive edges on an individual cell. Micropatterns were made on a layer of deformable poly-acrylamide gel, in which beads were incorporated (Beningo et al., 2002; Vignaud et al., 2014). From the tracking of bead displacements, the forces exerted by the cells were mapped, and the total strain energy the cells transmitted to the substrate was computed (Butler et al., 2002; Martiel et al., 2015). Strikingly, E/M cells displayed lower traction forces than E cells, whereas M cells displayed higher traction forces (Figure 4c). This outcome appeared independent of the geometrical configuration of the micropattern. Using a rectangle micropattern, making an HMEC adopt a shape reminiscent of that adopted in tissue, the same relationship was observed, i.e. cells in the E/M state were much less contractile than those in the two other states (Figure 4d). Importantly, this showed that some of the biophysical properties of cells in E/M states are not between those of E and M cells, but beyond them.

Second, we evaluated HMECs on H micropatterns, after brief exposure to compounds that disrupted (blebbistatin or Y27632, 6 hours) or stimulated acto-myosin activity (calyculin A, 15 minutes; Figure 5). Given that M cells had displayed high traction forces, they were treated with blebbistatin or Y27632, and both treatments resulted in the polarity indices shifting from negative to positive (Figure 5a-c). This transition in the polarity indices was due to the positioning of the nuclei being more distal from the intercellular junction. Conversely, given that E/M-cells had displayed low traction forces, they were incubated with calyculin A, and this resulted in the polarity index shifting from near zero to negative, due to the positioning of the nuclei more proximal the intercellular junction (Figure 5d-f).

**Figure 5:**
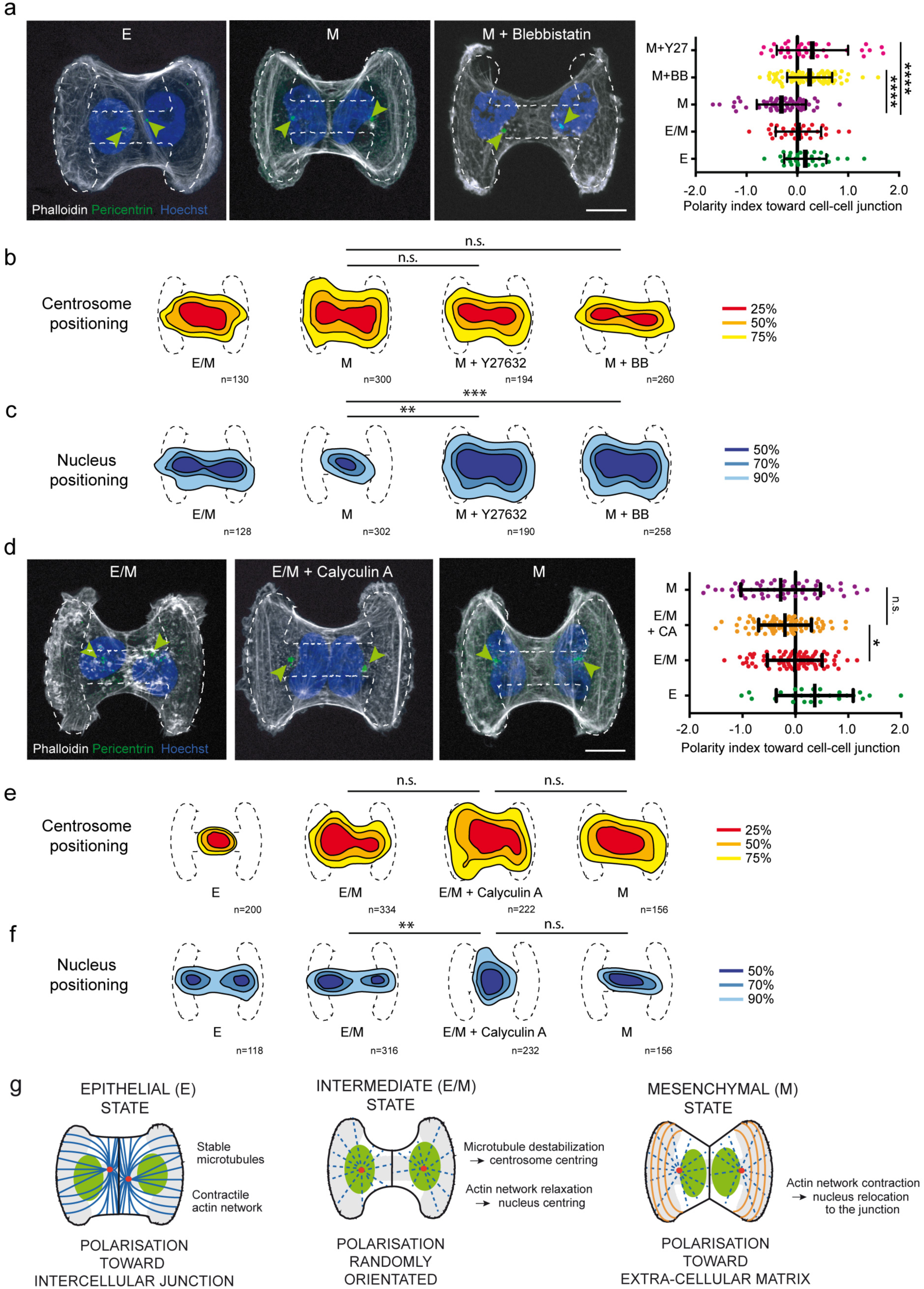
Acto-myosin contraction regulates nucleus positioning and polarity reversal in M cells. **a** Images show F-actin (white), centrosome (green) and nucleus (blue) in cells in the epithelial (E) state, epithelial/mesenchyme (E/M) state and M state treated with 25 µM blebbistatin. Scale bar is 20 µm. The graph shows the quantification of cell polarity indices on H micropatterns in E, E/M, M cells, and in M cells treated with blebbistatin (25 µM) or Y27632 (20 µM) for 6 hours. ****: p<0.0001 by one-way ANOVA and Tukey’s multiple comparison test. n>32, N=3 independent experiments. Error bars indicate SD. **b** Centrosome density maps representing the regions in which 25% (red) and 75% (yellow) of centrosomes in cell doublets can be found in each condition. n.s.: no significant difference. **c** Nucleus density maps representing the regions in which 50% (dense blue) and 90% (light blue) of nuclei in cell doublets can be found in each condition. **: p<0.01, ***: p<0.001 from two-sample kernel density-based test, n>128 cells. **d** Images show F-actin (white), centrosome (green) and nucleus (blue) in E/M cells, in E/M cells treated with 1 nM calyculin A for 15 minutes, and in M cells. The graph shows a quantification of polarity indices in each condition *: p<0.05 by one-way ANOVA and Tukey’s multiple comparison test. Error bars indicate SD. n>28, N=3 independent experiments. **e** Centrosome density maps representing the regions in which 25% (red) and 75% (yellow) of centrosomes in cell doublets can be found in each condition. n.s.: no significant difference. **f** Nucleus density maps representing the regions in which 50% (dense blue) and 90% (light blue) of nuclei in cell doublets can be found in each condition. **: p<0.01 from two-sample kernel density-based test, n>118 cells. **g** Schematic representation of cell polarity as a biomarker of EMT states. E cells have a stable microtubule network with contractile actin directing cell polarity (nucleus-centrosome axis) toward cell-cell junction. E/M cells have less stable microtubules and a low actin contractility leading to a random orientation of cell polarity. M cells also have less stable microtubules than E cells but high acto-myosin contractility, which directs cell polarity towards the extracellular matrix.

Interestingly, neither the relaxation of acto-myosin activity in M cells nor its stimulation in E/M cells had significant impacts on centrosome positioning (Figure 5b and 5e). Rather, the modulation of acto-myosin activity affected cell polarity by acting specifically on the positioning of the nucleus (Figure 5c and 5f and Supplementary Figure S3a). Similar results could be obtained by constitutively activating acto-myosin activity in E/M cells with the expression of an active form of RhoA (Supplementary Figure S3b and S3c) or by relaxing acto-myosin activity in M cells by plating them on soft substrates (Supplementary Figure S3d and S3e).

Altogether, these results pointed at key sequential roles of the remodeling of cytoskeleton networks during the intracellular reorganization that accompanies the reprogramming of cell states during EMT. In the E state, the cell is cohesive, with an off centered centrosome proximal to the intercellular junction (Figure 5g). In the E/M state, cell-cell cohesion is low, manifested by a smaller intercellular junction and the destabilization of microtubules, leading to the repositioning of the centrosome at the cell center. The cell is also poorly contractile, manifested by the central positioning of its nucleus, and thus an overall random orientation of its nucleus-centrosome axis (Figure 5g). In the M-state, cell-cell cohesion is also low with the centrosome position at the cell center. However, the cell is also highly contractile, manifested by the nucleus repositioned distal from cell-matrix adhesion towards the cell-cell junction, and by a preferential orientation of the nucleus-centrosome axis towards the extracellular matrix (Figure 5g).

These observations on cytoskeleton networks were made possible by confining cells in highly controlled spatial conditions imposed by defined stereotypic adhesion patterns that prevented cell motility. To further investigate and quantify the role of intercellular adhesion, polarity and contractility on a cell’s motility, we used different assays in which the constraints on a cell’s motility were relaxed.

### Selective increase of cell migration in E/M cells

First, a quantitative cell-scatter assay was used to evaluate cell-cell cohesion (Figure 6a). Individual cells were plated on 300 micron-long fibronectin-coated bars. A day later, cells had divided and the distance between daughter-cell nuclei was measured (Figure 6b). E cells remained attached and thus displayed short inter-nuclei distances (Figure 6c). By contrast the expection was confirmed that cells with a reduced capacity for cell-cell cohesion could scatter and this would be manifested by longer inter-nuclear distances. Hence both E/M and M cells tended to separate more frequently than E cells (Figure 6c). By way of controls, for the parental HMEC line, doxycycline had no effect increasing the frequency of daughter cells separating, whereas a 5-day treatment with TGFβ increased that frequency (Supplementary Figure S2d).

**Figure 6:**
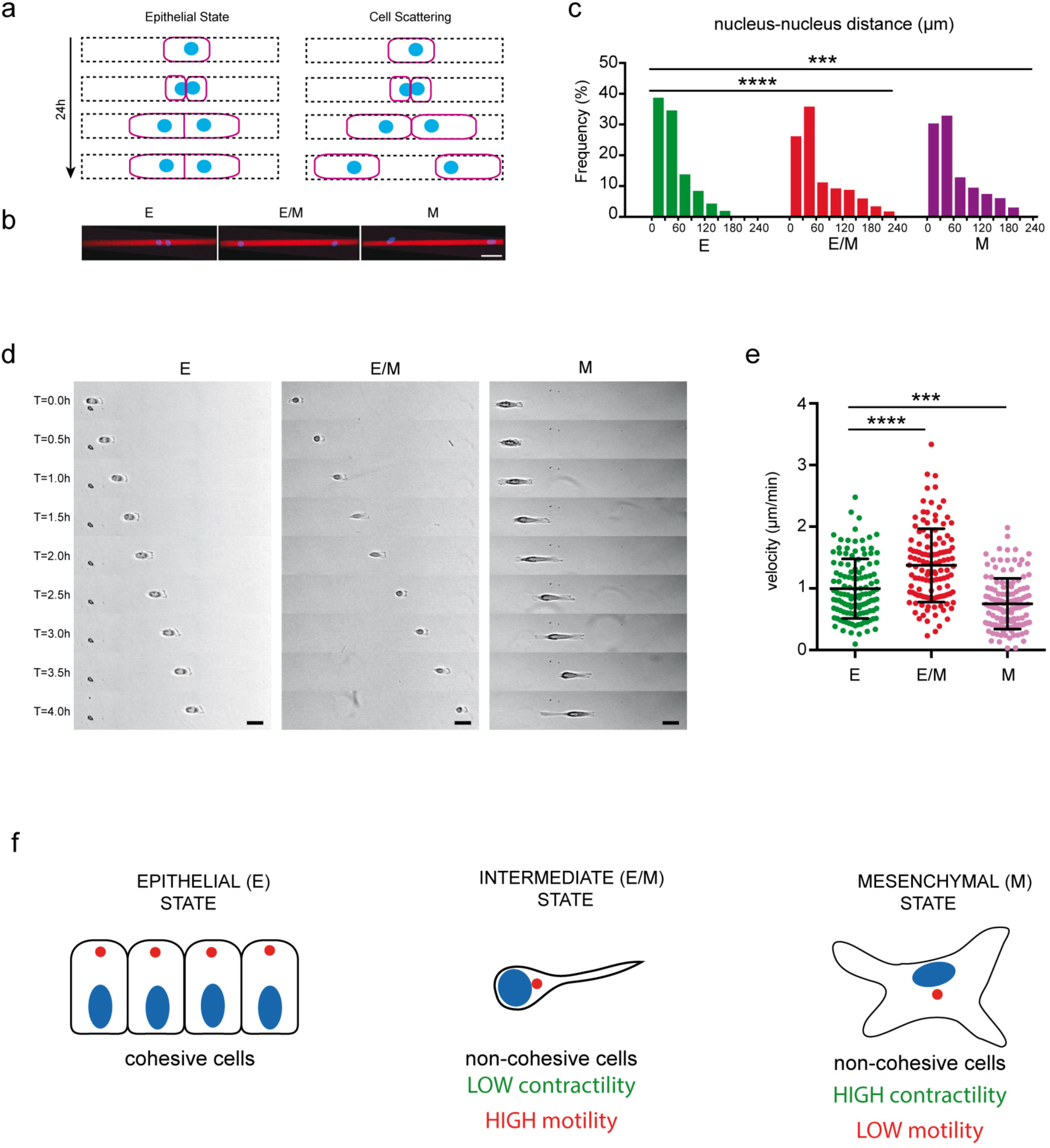
Scattering and migration of E, E/M and M cells. **a** Schematic representation of the cell-scattering assay with a single end point at 24 hours. Thin 300 µm long rectangular micropatterns initially containing a single cell are considered after 24 hours incubation to allow cells to divide once. The distance between the nuclei of the two daughter cells is measured as a proxy for their cohesiveness. **b** Images showing nucleus (blue) location on micropatterned lines (red) in cell-scattering assays. **c** Quantification of nucleus-nucleus distance in cell doublets in cell scattering assays. ***: p<0.001, ****: p<0.0001, by Kruskal-Wallis multiple comparison test. Minimum of n=589, N=3. **d** Images show sequences of images acquired in transmitted light during a single-cell migration assay of 4 hours. E, E/M and M cells were plated on 10 µm wide micropatterned lines and recorded in video-microscopy. Scales bar represents 20 µm. **e** Quantification of single-cell velocity on micropatterned lines. ***: p<0.001, ****: p<0.0001, by One-Way ANOVA test and Tukey’s multiple comparison. Minimum of n=109, error bars indicate SD, N=2 independent experiments. **f** Schematic representation of the biphasic changes in biophysical properties of cells during EMT progression. When cells pass from E to E/M and to M states, rather than a gradual continuum, contractility and motility phenotypes undergo biphasic changes: the contractility is first low then high, whereas motility (capacity) was first high then low. Hence the intermediate (E/M) state can be characterized as having the lowest contractility and the highest motile potential.

Second, a single cell motility assay was used to characterize cell-intrinsic migration capacities, in which cell velocities along 15-micron-wide lines of fibronectin were calculated. This assay is considered to better recapitulate 3D migration than classic 2D-migration assays (Doyle et al., 2009)(Figure 6d). Cell migration is inversely related to the tensional state in the cell. High traction forces are associated with extensive cell spreading, as well as the assembly of large focal adhesions, impeding the displacement of cell body. Conversely, a poorly contractile cell spread less, and detaches more easily, which permits a higher motile velocity (Leal-Egaña et al., 2017). Consistent with the observed levels of cell traction forces (Figure 4c,d) E/M cells migrated faster than E cells, whereas M cells migrated slower (Figure 6e). Again, by way of controls, for the parental HMEC line, doxycycline had no effect and the 5-day treatment with TGFβ also lowered cell migration velocity (Supplementary Figure S2e).

Interestingly, these results revealed that the mechanical strength and migration capacities of E/M and M cells were polar opposites with respect to those capacities in E cells, indicating that the E/M state has distinct and specific biophysical properties that are not intermediate between the E and M states. E/M cells displayed lower traction forces and thus higher migration velocities than E and M cells (Figure 6f). This observation supports the emerging concept that invasive tumor cells are in the E/M state rather than the M state. We further tested this idea by evaluating the characteristics of well-established invasive breast cancer cell lines with respect to the structural and mechanical readouts used on HMECs.

### Triple-negative breast cancer cells displayed the structural features of E/M state

Triple negative breast cancer cells (TNBCs) are resistant to hormone treatment and thus associated with poor prognosis (Grigoriadis et al., 2012). TNBCs are also prone to induce the formation of metastases (Dent et al., 2009; Hudis and Gianni, 2011). To compare their structural and mechanical properties to those of the E, E/M and M states of HMEC cells, several TNBC lines were cultured as cell doublets on H-shaped fibronectin-coated micropatterns over 24 hours (Figure 7a). The BT-20 and HCC38, TNBC cell lines, considered as low-invasive (Chavez et al., 2010; Grigoriadis et al., 2012), had polarity indices similar to non-transformed epithelial MCF10A cells and HMEC in the E state (Figure 7b and Figure 1g). Most other TNBC cell lines (MDA-MB-157, MDA-MB-231, MDA-MB-436 and HCC1937), displayed almost random orientation of their nucleus-centrosome axis with a polarity index close to zero similar to HMEC in the E/M state (Figure 7b). In those cells, the loss of polarity was associated with both nuclear and centrosome positioning at the cell center (Supplementary figure S4). Notably, none of the TNBC cell lines displayed the complete reversal of cell polarity that is typical of M states.

**Figure 7:**
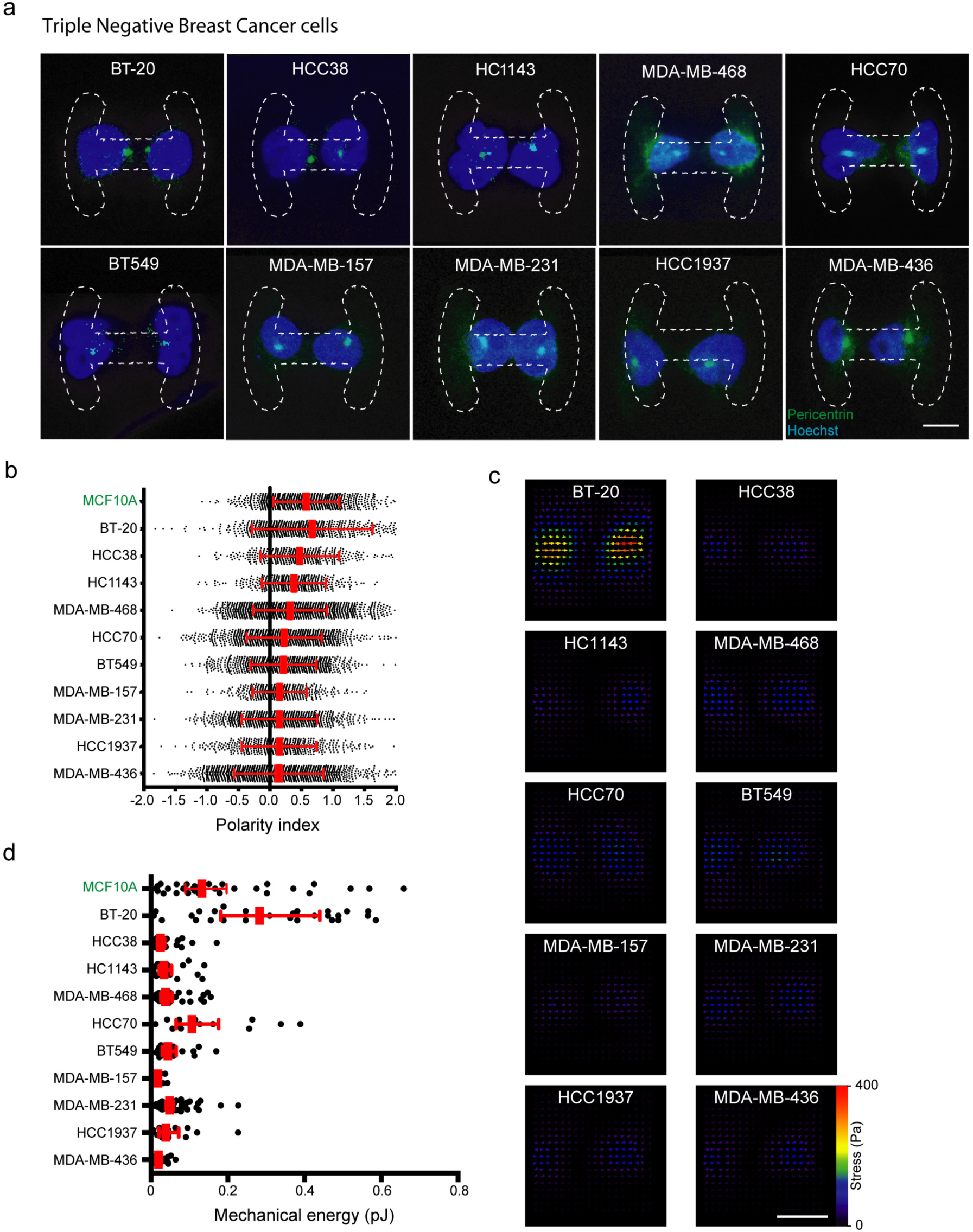
Cell polarity and contractility of Triple Negative Breast Cancer cells (TBNCs). **a** Cell-polarity in a panel of TNBC lines. Representative images, from maximum projections of Z-stacks, showing cell doublets cultured during 24 hours on H-shaped micropatterns (drawn with dashed lines). Cells were stained with pericentrin (green) and Hoechst (blue) for centrosome and nucleus location, respectively. Scale bar represents 20 μm. **b** Polarity indices in TNBC lines ranked by average polarity, and in comparison with MCF10A human mammary epithelial cell line. Error bars indicate SD. n>427 for each cell line, N=3 independent experiments. **c** Averaged traction stress field (in Pascal) in TNBC lines cultured on rectangle micropatterned polyacrylamide hydrogels. N>2 independent experiments. **d** Mechanical energy of cell traction of TNBC cell lines, and compared with MCF10A cell line. n>14 for each cell line, error bars indicate 95% confidence interval, N>2 independent experiments.

Given that the low magnitude of traction forces was a clear signature of E/M states in HMECs, we plated TNBCs on micropatterned poly-acrylamide gel to evaluate the tensional properties of these cells. Strikingly, all TNBCs, apart from BT-20, displayed very low levels of contractile energy, similar to HMECs in E/M state and significantly lower than MCF10A or HMEC in E or M state (Figure 7c, d). These results showed that with regard to intracellular organization and tensional states, TNBC lines shared the same phenotype as HMECs in E/M state.

## Discussion

Using cell models of intermediate and complete EMT, respectively induced through the expression of a single EMT-TF or through the treatement by TGFβ, we were able to shed light on the intermediate states of EMT. These intermediate states have been previously characterized by specific protein-expression profiles and tissue localization (Krebs et al., 2017; Kröger et al., 2019; Morel et al., 2017; Pastushenko et al., 2018). Here, we investigated cell internal organization and key organelles positioning during the intermediate states of EMT. Our results indicate that intermediate states can be characterized by specific architectural and mechanical parameters: central position of the centrosome, central position of the nucleus and low acto-myosin contractility.

Interestingly, based on the structural and mechanical readouts we defined, the panel of TNBC cell lines had phenotypes corresponding to intermediate (E/M) states of EMT, closer to that of HMEC over-expressing ZEB1 than HMECs exposed to TGFβ (the M state). These phenotypic similarities between ZEB1-expressing HMECs and TNBCs echoes previous observations in 3D cultures and in tissues (Spaderna et al., 2008) showing that the expression of ZEB1 is sufficient to induce invasion, resistance to anoikis and resistance to drug treatments (Caramel et al., 2018; Kröger et al., 2019), as is the case for TNBCs. These similarities were also consistent with a recent study on TNBCs showing that the expression of ZEB1, and not the expression of advanced mesenchymal markers, was specifically correlated with poor clinical outcome (Jang et al., 2015). Interestingly, the stimulation with TGFβ induced a distinct phenotype, with a strong contractility and clear polarization toward the extracellular matrix, even though the TGFβ pathways proceed via the induction of ZEB1 (Tsubakihara and Moustakas, 2018). We interpreted the differences between the E/M-state and M-state in our experimental models as the consequence of the activation of the RhoA pathway, which is also downstream of TGFβ (Bhowmick et al., 2001; Kardassis et al., 2009; Tavares et al., 2006) (Figure 5 and Supplementary figure S3).

The early stages of EMT were associated with centrosome repositioning to the cell center in our cell models. In epithelial cells, the centrosome is off-centered and positioned proximal to the intercellular junction (Burute et al., 2017; Rodriguez-Fraticelli et al., 2012). Similar off-centering has been observed in different conditions and the common underlying mechanism is considered to involve the stabilization of microtubules proximal to the intercellular junction and the capture and pulling of the centrosome towards the junction by those microtubules (Combs et al., 2006; Schmoranzer et al., 2009; Sipe et al., 2013). Indeed, microtubules and junctional adhesions mutually reinforce their stabilization (reviewed in (Vasileva and Citi, 2018)) (Chausovsky et al., 2000; Huang et al., 2011; Ligon et al., 2001; Meng et al., 2008; Shahbazi et al., 2013; Stehbens et al., 2006). Here we found that in early stages of EMT, microtubules were destabilized and the centrosome moved away from the junction. This interpretation is consistent with our previous observation that Par3 is downregulated at the junction between MCF10A cells with TGFβ, and that the overexpression of Par3 can overide the effect of TGFβ and maintain the centrosome at the junction (Burute et al., 2017). As such, the centrosome displacement away from the intercellular junction in E/M cells is likely to reflect the increased fragility of the intercellular connection between the cells that have initiated EMT. Similar crosstalk between the centrosome and intercellular junction are associated with cell scattering and EMT in development processes in living organisms (Bornens, 2018). Neuronal-cell delamination and migration from the neural tube involves an increase in microtubule dynamics and centrosome displacement away from junction with neuronal-plate cells (Das and Storey, 2014; Kasioulis et al., 2017). Consistent with observation that microtubule destabilization occurs in the early stages of EMT, TNBCs can be directed towards an epithelial state by the known microtubule stabilizer eribulin (Yoshida et al., 2014). Thus, our study establishes centrosome positioning as a potential key marker for identifying individual cells in early stages of EMT in breast cancer.

The initiation of EMT and establishment of the intermediate E/M state was characterized by a significant reduction in acto-myosin contractility in our cell models. However, this contractility was recovered or even enhanced when the cell transitioned from the E/M-state to the M state. This biphasic change in contractility contrasts with the view of EMT being a progressive and monotonic transition, in which parameters either increase, or decrease, from the E state to the M state. In addition, we found that E/M cells had higher migratory properties than M cells, which is consistent with recent in vivo data showing that E/M cells can be found at the invasive front of primary tumors (Pastushenko et al., 2018). The correlation between low contractility and fast migration is consistent with our previous observation that fast migrating cells generate low traction forces (Leal-Egaña et al., 2017). The low-contractile-force phenotype differs from the view that contractile tension increases with cell transformation in cancer (Paszek et al., 2005) and that a higher rigidity of the extracellular environment and a high-contractile-force phenotype favor cell invasiveness (Butcher et al., 2009; Kraning-Rush et al., 2012; Mierke et al., 2011). Rather, our results suggest that the relaxation of contractile forces is associated with cell scattering and migration, as it has been observed in HMECs with super-numerary centrosomes that form invasive structures in 3D cultures (Godinho et al., 2014), in cells at the front of invasive colorectal carcinomas (Libanje et al., 2019) or in metastatic osteosarcoma cells (Holenstein et al., 2019). By contrast, a high-contractile-force phenotype is more likely to promote primary tumor growth, cell survival in circulation, and at a secondary site, cell adhesion, aggregation, and the formation of a metastatic tumor (Rodriguez-Hernandez et al., 2016; Tavares et al., 2017). Hence, both low- and high-contractile-force phenotypes are central for cancer progression. Therefore the characterization of the contractile force phenotype of tumor cells in controlled conditions, i.e. in a given cell type with specific EMT inducers, would help to clarify some conflicting interpretations on the role of certain tumorigenic factors.

Intermediate states of EMT have also been associated increased tumor-cell plasticity, or stemness, i.e. the capacity of a tumor cell to differentiate into multiple lineages, a capacity which may favor survival of the line because of the greater chance that certain progeny metastasize and form secondary tumors (Brabletz, 2012; Puisieux et al., 2018). Therefore the low-contractile-force phenotype of E/M cells may also be correlated with stemness. This is suppoted by the observations that the down-regulation of CK2β in mammary epithelial cells induces the expression of stemness markers (Duchemin-Pelletier et al., 2017), mechanical relaxation (Tseng et al., 2011) and the acquisition of high migration velocities (Leal-Egaña et al., 2017). This correlation between stemness and the low-contractile-force phenotype also echoes the widely-used Rho kinase inhibitor Y27632 to maintain the undifferentiated growth of pluripotent stem cells in culture in culture media; an inhibitor which is also potently relaxes acto-myosin contractility (Gauthaman et al., 2010; Pakzad et al., 2010). Therefore in tumor cells in intermediate states of EMT, a causal relationship between stemness and the low-contractile-force phenotype is an intriguing possibility that deserves further investigations.

## Materials and Methods

### Cell culture, plasmids, transfection and drug treatment

BT20, MDA-MB-157, MDA-MB-231, MDA-MB-436 and MDA-MB-468 cell lines were cultured in Dulbecco’s Modified Eagle Medium (31966, Gibco) supplemented with 10% FBS (50900, Biowest) and 1% antibiotic-antimycotic (15240-062, Gibco). BT-549, HCC38, HCC70, HCC1143, HCC1937 were cultured in RPMI 1640 (61870, Gibco) with 10% FBS and 1% antibiotic-antimycotic. MCF10A were cultured in MEGM growth medium (CC3151, Lonza) supplemented with 100 ng/ml cholera toxin (C8052, Sigma) according to ATCC protocol. HMEC-hTert-puro-Ptripz-ZEB1 and HMEC-hTert-puro Ptripz were cultured in MEM/F12 (31331, Gibco), 10% FBS (deprived of tetracycline, S181T, Biowest)+ 1% antibiotic-antimycotic + Insuline (10µg/ml, CC4136, Lonza) + hEGF (10ng/ml, CC4136, Lonza) + hydrocortisone (0.5µg/ml, CC4136, Lonza) + puromycin (0.5µg/ml, A1113803, Thermo Fischer Scientific). ZEB1 expression was induced by adding 1 µg/ml doxycycline (D3447, Sigma) to culture medium during 24hrs. EMT was induced by 5 ng/ml TGFβ1 (240-B-002, R&D Systems) treatment for up to 5 days. ROCK inhibition was achieved using 20 µM Y-27632 (ab120129, Abcam) and actomyosin contractility by 25 µM blebbistatin (B0560, Sigma) for 6 hours. Cell contractility was increased using 1 nM calyculin A (ab141784, Abcam) treatment for 15 minutes. pCS-6xMyc-RhoAG14V was kindly provided by Jean-François Côté (IRCM, Montréal) and transfected using lipofectamine 2000 (11668019, Thermo Fischer Scientific) in Opti-MEM (11058, Gibco) according to the procedure from the manufacturer. For culture on micropatterned coverslips, after trypsin treatment (12605, Gibco) cells were seeded at a density of 500000 cells per chip in culture medium for 24 hours incubation, except for the single-cell migration assay on line micropatterns: 10000 per chip. The cold-treatment assay was performed on plain coverslips after a seeding of 200000 cells per chip, cells were allowed to spread for 4 hours and then incubated or not on ice for 15 minutes prior to fixation. All live cells were incubated and imaged in a humidified environment at 37°C with 5% CO_2_.

### Antibodies and cell staining

Primary antibodies used for immunostaining included anti-pericentrin (ab4448, abcam, 1/1000 dilution), anti-c-myc 9E10 (MM4439, Sigma, 1/1000 dilution), anti-E-cadherin (610181, BD, 1/500 dilution) and anti-α-tubulin (MCA776, ABD Serotech, 1/500 dilution). Anti-rabbit-Alexa488 (A27034, Thermo Fischer Scientific) and anti-rat-Alexa594 (A21471, Thermo Fischer Scientific) were used as secondary antibodies at 1/500 dilution. F-actin staining was performed using phalloidin-Alexa594 (A12381, Invitrogen, 1/500 dilution) or phalloidin-ATTO488 (49409, Sigma, 1/500 dilution) and nuclei were stained with Hoechst 33342 (H1399, Thermo Fischer Scientific). For immunostaining, cells were fixed for 10 minutes in cold methanol (−20°C, 34860, Sigma) followed by 3 washing steps in PBS (18912, Gibco). Cells were permeabilized in 0,2% triton X100 (X100, Sigma) for 15 minutes at room temperature, then blocking was performed for 10 minutes at room temperature in 3% BSA (A4737, Sigma). Cells were incubated with primary antibody in 3% BSA for 45 minutes at room temperature and with secondary antibodies in 3% BSA for 30 minutes at room temperature. After 3 PBS washes, chips were mounted on microscope slides using ProLong Gold antifade reagent with DAPI (Thermo Fischer Scientific). Live observation of microtubule and actin networks was performed after 500 nM SiR-Tubulin (SC002, Tebu-bio) and SiR-Actin (SC001, Tebu-bio) staining for 3 hours, respectively. For immunoblotting, anti-ZEB1 (HPA027524, Sigma, 1/1000 dilution), anti-ZEB2 (HPA003456, Sigma, 1/500 dilution), anti-SNAIL+SLUG (ab180714, abcam, 1/500 dilution), anti-E-cadherin (SC8426, Santa-Cruz, 1/5000 dilution), anti-vimentin (MA5-14564, Thermo Fischer Scientific, 1/1000 dilution), and anti-GAPDH-HRP (SC47724, Santa-Cruz, 1/200 dilution), anti-mouse-HRP (SA-1-100, Thermo Fischer Scientific, 1/5000 dilution), and anti-rabbit-HRP (800-367-5296, Jackson ImmunoResearch, 1/5000 dilution) were diluted in 1% Tween-PBS with 3% BSA.

### RT-qPCR assay

Real-time PCR intron-spanning assays were designed using the Universal Probe Library Assay-design Centre software (Roche Applied Science). RNA was prepared using an extraction column [RNeasy mini Kit (Qiagen) for HME-derived cells; RNeasy micro-Kit (Qiagen) for sorted cells] according to manufacturer’s instructions. For primary cells, a whole genome pre-amplification (Amplification kit system ovation QPCR (Nugen)) has been assessed on mRNA. One microgram of total RNA was used for cDNA synthesis (DyNAmo cDNA synthesis, Thermo-scientific), and quantitative PCR (qPCR) analysis was performed with 3 ?l of a 1/10 dilution of resulting cDNAs. DNA amplification was monitored by real-time PCR using a CFX96 (Bio-Rad) and analyzed with the Bio-Rad CFX manager software. The relative quantification of gene expression was performed using the comparative CT method, with normalization of the target gene to one of the two endogenous housekeeping genes, namely 36B4 or HPRT1.

List of primer sequences used for Q-PCR analysis: Human ZEB1 AGG GCA CAC CAG AAG CCA G and GAG GTA AAG CGT TTA TAG CCT CTA TCA, human ZEB2 aag cca ggg aca gat cag c and gcc aca ctc tgt gca ttt ga, human SNAI1 GCT GCA GGA CTC TAA TCC AGA and ATC TCC GGA GGT GGG ATG, human CDH1 ccc ggg aca acg ttt att ac and gct ggc tca agt caa agt cc, human VIM gac cag cta acc aac gac aaa and gaa gca tct cct cct gca at. »

### Cell micropatterning and microfabrication

H-shape fibronectin coated cell-adhesive micropatterns were provided from CYTOO (www.cytoo.com), as H 1100 µm^2^, in 20 mm^2^ chip format. Other micropatterns were fabricated according to (Azioune et al., 2010). Glass coverslips, coated with 0.1 mg/ml (PLL)-poly-ethylene-glycol (PLL20K-G35-PEG2K, JenKem) in 10 mM HEPES at pH 7.4 for 30min, were oxidized through oxygen plasma (FEMTO, Diener Electronic) for 10 seconds at 30 W before exposure to 165 nm UV (UVO-Cleaner, Jelight) through a chromium photomask (Toppan) for 5 minutes. Coverslips were incubated with 20 mg/ml of FN (FF1141, Sigma) and 20µg/ml fluorescently labelled fibrinogen (Alexa Fluor 647 conjugate, F35200, Thermo Fischer Scientific) in 100 mM sodium bicarbonate (144-55-8, Sigma) solution at pH 8.3 for 30 minutes.

Soft micropatterns were prepared by polymerizing a mix of acrylamide (A9099, Sigma) and bis-acrylamide (294381, Sigma) in a respective ratio of 8%/0.264%, on a micropattern glass coverslip to allow protein transfer to hydrogel as in (Vignaud et al., 2014). Polyacrylamide solution was de-gassed for around 30min and mixed with passivated fluorescent beads (red fluorescent FluoSpheres carboxylate-modified microspheres, 0.2µm F8810, Thermo Fischer Scientific) by sonication before adding ammonium persulfate and tetramethylethylenediamine. Polymerization was achieved on the photomask in a period of 21 minutes under a silanized coverslip. The silanized coverslip was removed in the presence of sodium bicarbonate and the gels stored overnight at 4°C in sodium bicarbonate. Coverslips were washed in sterile PBS (14190, Gibco) before cell seeding and mounting in a magnetic chamber (Chamlide, LCI) for TFM processing.

### Image acquisition

#### Cell polarity and scattering

Images were obtained from 3D acquisitions using an Eclipse Ti-E inverted microscope equipped with a CSUX1-A1 Yokogawa confocal head, an Evolve EMCCD camera (Roper Scientific, Princeton Instrument). The system was controlled by MetaMorph software (Molecular Devices). For cell polarity and scattering assay, Nikon Plan-APO 40x-0.95 and Plan-APO VC 20x-0.75 dry objectives were used respectively. Z-maximum projections were achieved.

#### Cytoskeleton architecture

F-actin and microtubule networks were imaged in 3D in fixed cells using a Zeiss LSM880 confocal laser-scanning microscope in Airyscan mode, equipped with a Zeiss Plan-APO ×63–1.46 oil immersion objective. Airyscan reconstruction was achieved using the Zeiss basic plugin. Z-maximum projections were achieved. Microtubule fluorescence images were inverted in ImageJ (black to white, white to black) to appreciate the details of the network.

#### Live microscopy

Images were collected on a Zeiss LSM880 confocal laser-scanning microscope equipped with a Zeiss Plan-APO ×63–1.46 oil immersion objective. Overall cell displacement for measurement of single-cell–migration velocity was assessed from phase-contrast time-lapse acquisitions over 24 hours with 30minutes intervals. Remodeling of actin and microtubule cytoskeletons were imaged in live and in 3D over 20 minutes with 30 sec intervals. Z-maximum projections of microtubule network were achieved for display.

#### Traction force microscopy

Images of cells, beads and patterns were acquired with a confocal spinning disk system (Eclipse Ti-E Nikon) equipped with a CSUX1-A1 Yokogawa confocal head, an Evolve EMCCD camera (Roper Scientific, Princeton Instrument) and a Nikon CFI Plan-APO VC ×60–1.4 oil immersion objective).

#### Immunoblotting

HRP signal was revealed using Amersham Western Blotting detection reagents (RPN2209, GE Healthcare) according to the instruction of the manufacturer, and acquired using a Chemidoc Imaging System (Bio-Rad). Signal densities were quantified using ImageJ (NIH; http://rsb.info.nih.gov/ij).

### Image analysis

#### Cell polarity

Image processing was performed using an ImageJ macro. First, fluorescent cell-adhesive pattern were individualized using a template matching method. Second, nucleus and centrosome detection were realized based on threshold and size filtering within each cell of the doublet. Taking into account that only cell doublets were considered for further analysis, groups of cells encompassing two nuclei and two centrosomes were exclusively retained. The center of mass of the nucleus was computed and each centrosome was assigned to the closest nucleus. Third, the nucleus-centrosome and nucleus-nucleus vectors were computed. Then a post-processing step was realized in order to quantify the orientation and connectivity of each cell within a doublet. The inter-nuclear distance was measured as the length of the nucleus-nucleus vector. The cell-cell connectivity was defined as the difference between the area of the convex envelope surrounding the cell doublet and the area of the cell doublet.

#### Cell scattering

Nuclei were identified by thresholding using ImageJ. Images containing 2 nuclei only were considered. The distance between nuclei was computed also using ImageJ.

#### Single cell migration assay

Only single-cell displacement without any cell-cell contact during a period up to 4 hours was considered. Acquired images were analyzed using the manual tracking plugin of ImageJ to measure velocities.

#### Traction Force Microscopy

TFM and image analysis were carried out as described in (Martiel et al., 2015). All processing was carried out using ImageJ. Plugins and macros are available at https://sites.google.com/site/qingzongtseng/tfm. First, bead images were aligned to correct for experimental drift. Second, the displacement field was calculated using particle image velocimetry on the basis of a normalized cross-correlation algorithm following an iterative scheme. The final grid size for the displacement fields was 0.267 µm × 0.267 µm. The traction-force field was calculated by means of Fourier transformation of traction cytometry with a regularization parameter set to 2 × 10^−10^.

#### Organelle Mapping

Probabilistic density maps of nucleus and centrosome positioning were achieved according to (Duong et al., 2012; Schauer et al., 2010) based on ImageJ and R plugins. Density mapping was based on kernel density estimation. Briefly, raw images were segmented to structures of interest using ImageJ. Then a cell alignment was achieved from the normalized coordinates of the micropattern center by using R. Finally, kernel-density estimation was computed using R.

#### Actin flow measurement

From time-course 3D raw images, a time-dependent standard deviation was computed to reveal actin displacement by using ImageJ. After background subtraction, actin signal threshold was determined and the covered area calculated for each Z-section. Temporal color code representations were achieved from 50% height sections using ImageJ.

### Statistics and reproducibility

Statistical analysis was performed using GraphPad Prism software (Version 5.00, GraphPad Software). For each experiment, cell sampling and the number of independent replicates are indicated in figure legends. Data sets with normal distributions were analyzed with either the Student’s *t* test to compare two conditions, or with one-way ANOVA and Tukey tests to compare multiple conditions. Data sets that were not normally distributed in the normality test, or those data sets that should not be considered as having normal distributions, were analyzed with a Kruskal–Wallis test (multiple comparison). The comparison of a single pair of probabilistic density maps was performed using the two-sample kernel density-based test introduced in (Duong et al., 2012) by using R software. In the case of TFM and polarity-assay measurements, outlier identification and data cleaning were performed due to apparent defects in some individual measurements. Results are presented as mean and SD, or mean and 95% confidence interval as indicated in figure legends.

## Acknowledgements

This work was supported by grants from the European Research Council to Manuel Théry (ERC CoG 771599, ERC PoC 780458). Access to the imaging platform MuLife of DBSCI/IRIG was supported by funding from the Agence Nationale pour la Recherche (ANR-17-EURE-003). This work done by the team “EMT and cancer cell plasticity” led by Alain Puisieux was supported by la Ligue contre le Cancer, and the LabEx DEVweCAN. Tomoaki Nagai received a postdoc fellowship from Toyobo Biotechnology Foundation.

## Authors contribution

Yoran Margaron performed all experiments apart from the traction-force measurements on rectangular micropatterns, which were performed by Tomoaki Nagai. Laetitia Kurzawa helped on traction-force microscopy experiments. Laurent Guyon helped to analyze the data. Anne-Pierre Morel engineered and helped on the use of HMEC-ZEB1. Alice Pinheiro and Fabien Reyal provided and helped on the use of TNBCs. Laurent Blanchoin, Alain Puisieux, Yoran Margaron and Manuel Théry supervised the project. Yoran Margaron and Manuel Théry wrote the manuscript, which was further edited by all authors.

---

**Supplementary Figure S1:**
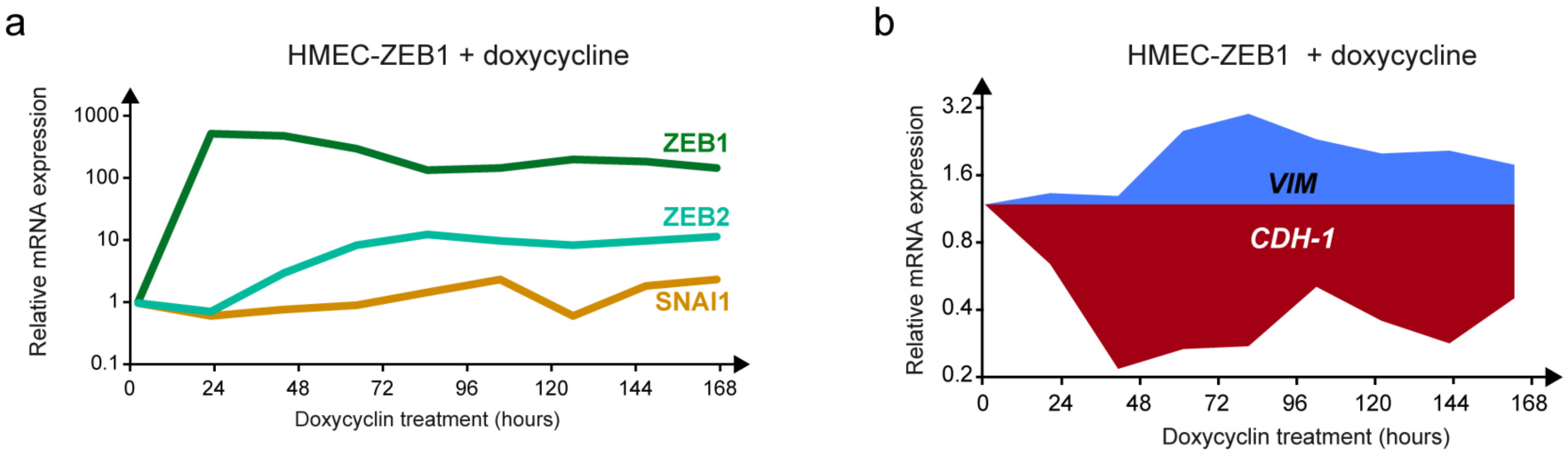
The HMEC-ZEB1 inducible system. **a** Graphs show ZEB1, ZEB2 and SNAI1 mRNA expression during doxycycline treatment over time in HMEC-ZEB1 inducible cell line. **b** Vimentin and E-cadherin mRNA expression measured by RT-qPCR during doxycycline treatment.

**Supplementary Figure S2:**
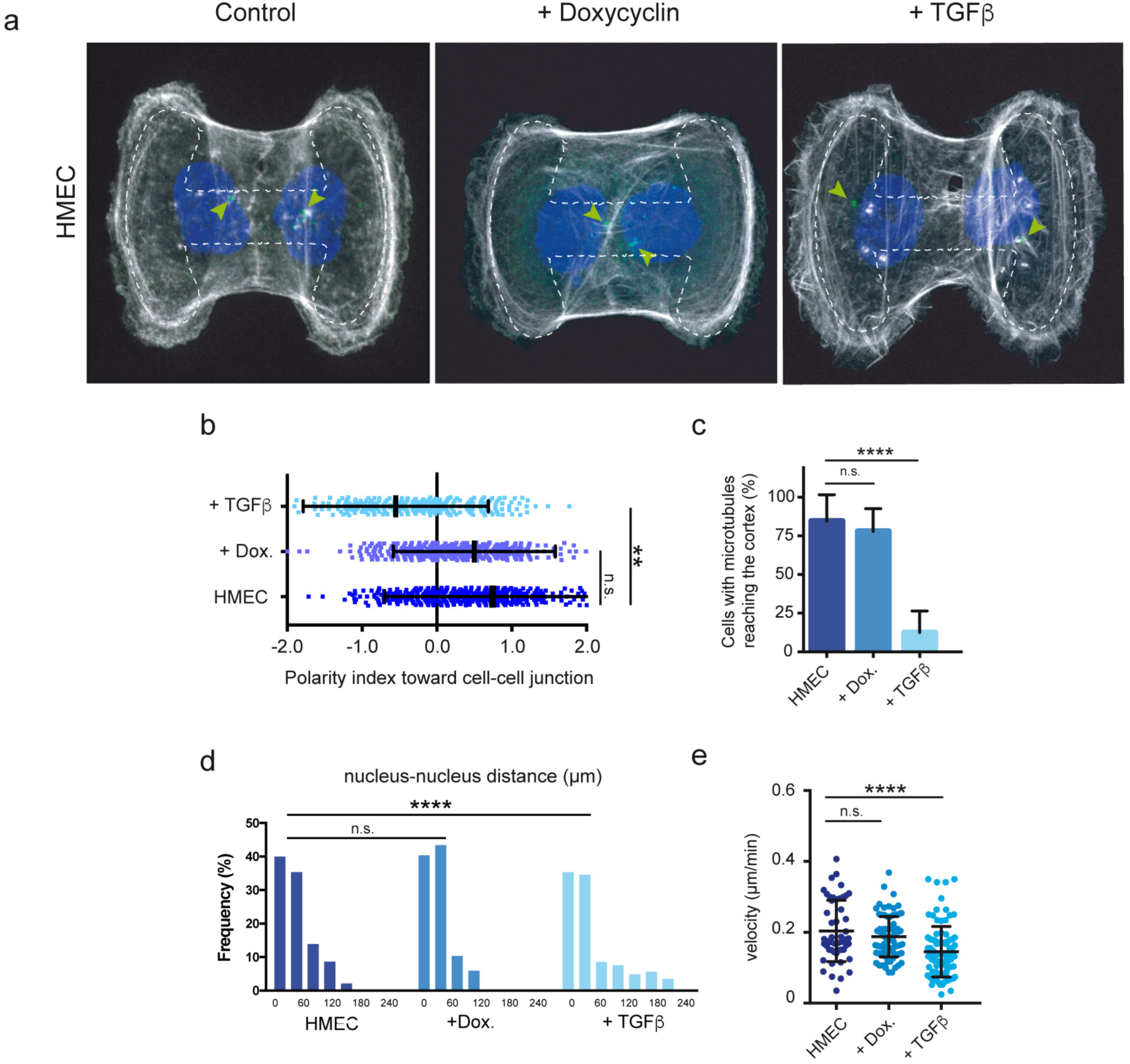
Effect of TGFβ and doxycycline treatments on the parental HMEC line. **a** Representative images showing HMEC doublets on H-shaped micropatterns (dashed lines) after being exposed to doxycycline for 24 hours, or TGFβ for 5 days. Cells were stained for F-actin (white), pericentrin (green) and Hoechst (blue). Arrowheads highlight centrosome positioning. Scale bar represents 20 µm. **b** Quantification of polarity indices in untreated HMECs, in cells treated with doxycycline for 24 hours, and in cells treated with TGFβ for 5 days.. **: p<0.01 by Student *t*-test. Error bars indicate SD, n>620 cells, N=3 independent experiments. **c** Quantification of the number of cells in which microtubules were identified at the cell cortex after cold treatment. ****: p<0.0001 by one-way ANOVA and Tukey’s multiple comparison test. Error bars indicate SD, n>50, N=3 independent experiments. **d** Quantification of nucleus-nucleus distance in cell-doublet scattering assay. ****: p<0.0001, by Kruskal-Wallis multiple comparison test. n>515, N=3 independent experiments. **e** Quantification of cell velocity in single-cell migration assay on 20 µm wide micropatterned lines in order to compare untreated HMECs with HMECs treated with doxycycline or TGFβ. ****: p<0.0001, by one-way ANOVA and Tukey’s multiple comparison test. Error bars indicate SD, n>50 N=2.

**Supplementary Figure S3:**
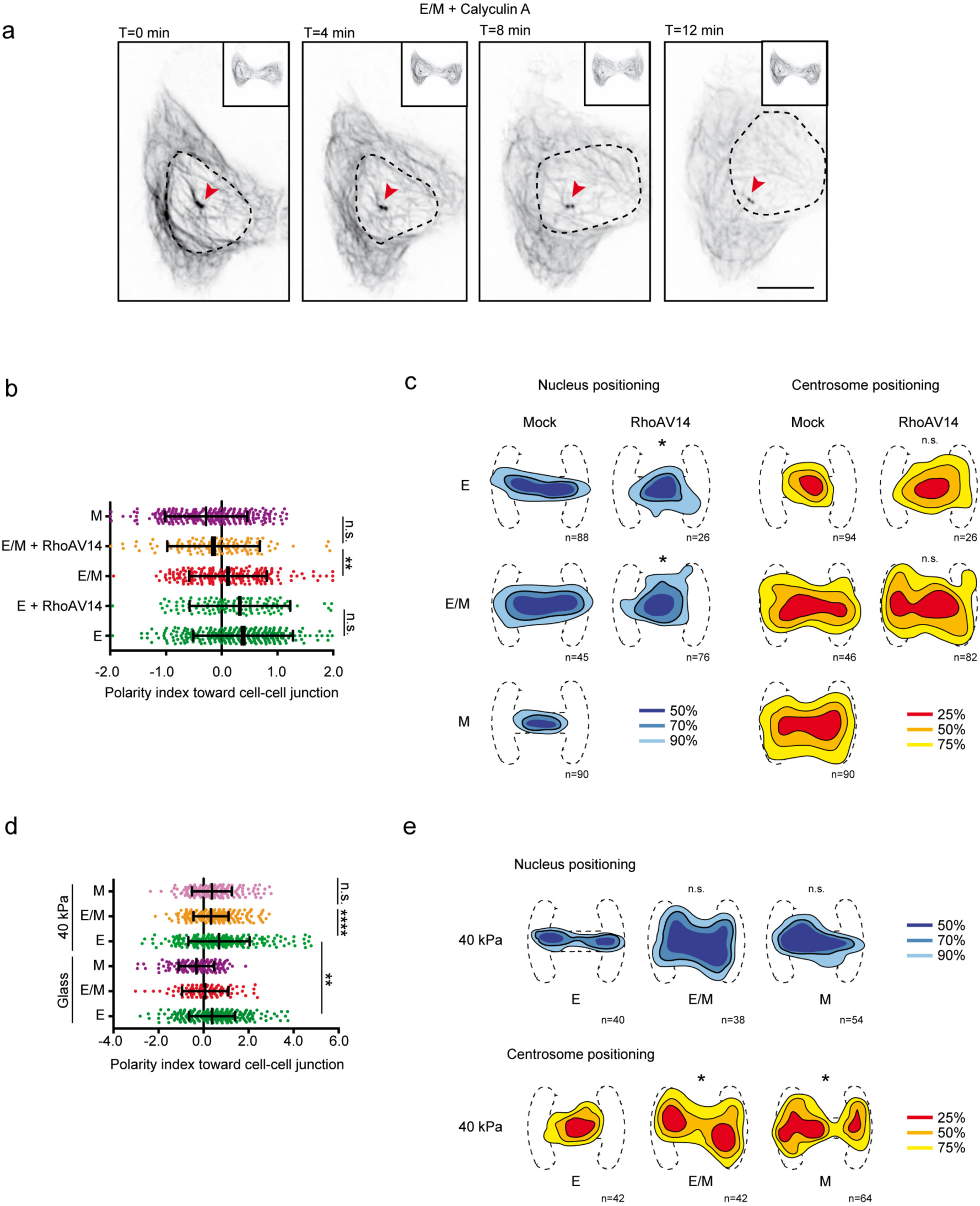
Impact of RhoA and substrate stiffness on nucleus positioning and cell polarity. **a** Sequence of representative maximal projection of Z-stacks from time-course experiment of at least 55 HMEC doublets stained for SiR-Tubulin (black) and Hoechst (dashed outlines), pretreated with doxycycline and cultured on H-micropatterns for 24 hours. Cell were treated with 1 nM calyculin A, and the images were acquired over 12 minutes by Airyscan microscopy. For clearer visibility, only one cell from the doublet is presented, right inset of each panel is the whole cell doublet. Red arrowheads show centrosome positioning. The nucleus is relocated toward the junction upon addition of calyculin A. Scale bar is 10 µm. **b** Quantification of polarity indices of HMECs containing a vector expressing RhoAV14 trangene or without transgene (mock control) in the epithelial (E) state, epithelial/mesenchyme (E/M) state, and in comparison with HMECs in the mesenchymal (M) state. ***: p<0.001, ****: p<0.0001, by Student *t*-test. Error bars indicate SD, n>124 cells, N=3. **c** Centrosome and nucleus density maps representing the regions in which 50% (dense blue) and 90% (light blue) of nuclei (left panel) and 25% (red) and 75% (yellow) of centrosomes (right panel) are found in cells described in (b). *: p<0.05, from two-sample kernel density-based test. **d** Quantification of polarity indices of E, E/M and M cells on plated on glass (control condition) and on soft (40kPa) micropatterned hydrogels. *: p<0.05, **: p<0.01, ****: p<0.0001, by Student *t*-test. Error bars indicate SD, n>96 cells, N=3 independent experiments. **e** Centrosome and nucleus density maps representing the regions in which 50% (dense blue) and 90% (light blue) of nuclei (left panel) and 25% (Red) and 75% (yellow) of centrosomes (right panel) are found in cells described in (d). *: p<0.05, from two-sample kernel density-based test.

**Supplementary Figure S4:**
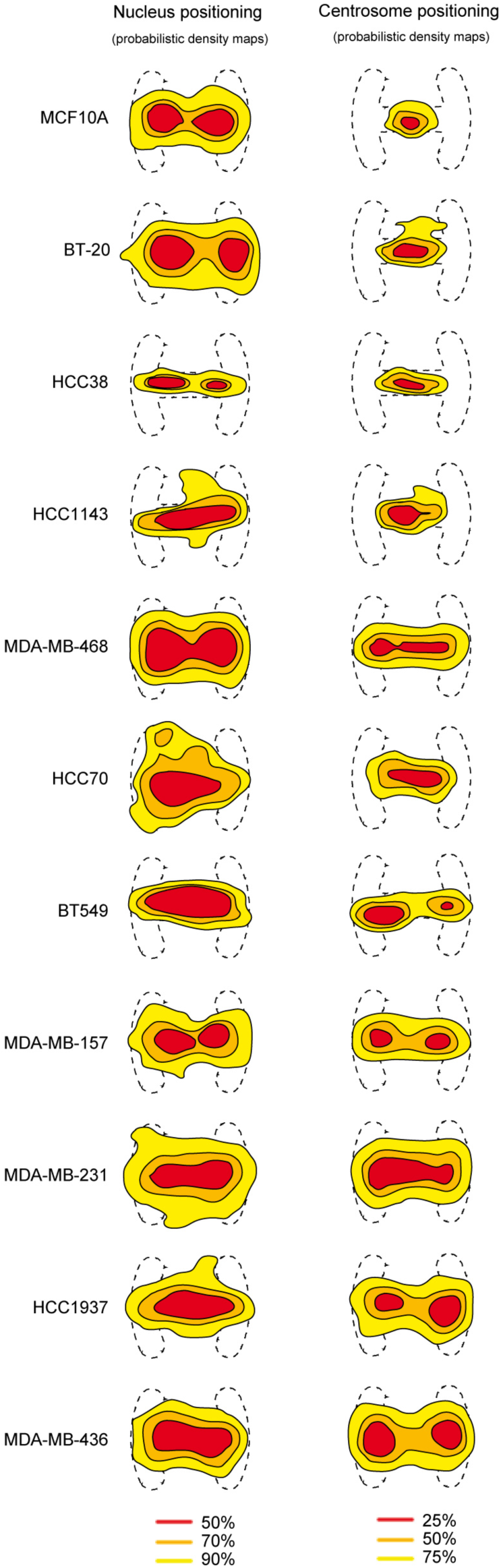
Probabilistic density maps of nucleus and centrosome positioning in TNBC cell lines. Maps representing the regions in which 50% (Red) and 90% (yellow) of nuclei and 25% (Red) and 75% (yellow) of centrosomes are found in a panel of TNBCs, and in comparison with MCF10A human mammary epithelial cell line.

## References

Aceto, N., Bardia, A., Miyamoto, D.T., Donaldson, M.C., Wittner, B.S., Spencer, J.A., Yu, M., Pely, A., Engstrom, A., Zhu, H., et al. (2014). Circulating tumor cell clusters are oligoclonal precursors of breast cancer metastasis. Cell 158, 1110–1122.

Alam, S.G., Lovett, D., Kim, D.I., Roux, K., Dickinson, R.B., and Lele, T.P. (2015). The nucleus is an intracellular propagator of tensile forces in NIH 3T3 fibroblasts. J. Cell Sci. 1901–1911.

Azioune, A., Carpi, N., Tseng, Q., Théry, M., and Piel, M. (2010). Protein Micropatterns. A Direct Printing Protocol Using Deep UVs.

Baccelli, I., Schneeweiss, A., Riethdorf, S., Stenzinger, A., Schillert, A., Vogel, V., Klein, C., Saini, M., Bëuerle, T., Wallwiener, M., et al. (2013). Identification of a population of blood circulating tumor cells from breast cancer patients that initiates metastasis in a xenograft assay. Nat. Biotechnol. 31, 539–544.

Beningo, K. a, Lo, C.-M., and Wang, Y.-L. (2002). Flexible polyacrylamide substrata for the analysis of mechanical interactions at cell-substratum adhesions. Methods Cell Biol. 69, 325–339.

Bhowmick, N.A., Ghiassi, M., Bakin, A., Aakre, M., Lundquist, C.A., Engel, M.E., Arteaga, C.L., and Moses, H.L. (2001). Transforming Growth Factor-β1 Mediates Epithelial to Mesenchymal Transdifferentiation through a RhoA-dependent Mechanism. Mol. Biol. Cell 12, 27–36.

Blanchoin, L., Boujemaa-Paterski, R., Sykes, C., and Plastino, J. (2014). Actin dynamics, architecture, and mechanics in cell motility. Physiol. Rev. 94, 235–263.

Bornens, M. (2008). Organelle positioning and cell polarity. Nat. Rev. Mol. Cell Biol. 9, 874–886.

Bornens, M. (2012). The Centrosome in Cells and Organisms. Science (80-.). 335, 422–426.

Bornens, M. (2018). Cell polarity: having and making sense of direction-on the evolutionary significance of the primary cilium/centrosome organ in Metazoa. Open Biol. 8, 180052.

Brabletz, T. (2012). To differentiate or not — routes towards metastasis. Nat. Rev. Cancer 12, 425–436.

Burute, M., and Thery, M. (2012). Spatial segregation between cell-cell and cell-matrix adhesions. Curr. Opin. Cell Biol. 24.

Burute, M., Prioux, M., Blin, G., Truchet, S., Letort, G., Tseng, Q., Bessy, T., Lowell, S., Young, J., Filhol, O., et al. (2017). Polarity Reversal by Centrosome Repositioning Primes Cell Scattering during Epithelial-to-Mesenchymal Transition. Dev. Cell 40, 168–184.

Butcher, D.T., Alliston, T., and Weaver, V.M. (2009). A tense situation?: forcing tumour progression. Nat. Rev. Cancer 9.

Butler, J.P., Tolić-Nørrelykke, I.M., Fabry, B., and Fredberg, J.J. (2002). Traction fields, moments, and strain energy that cells exert on their surroundings. Am. J. Physiol. Cell Physiol. 282, C595–605.

Caramel, J., Ligier, M., and Puisieux, A. (2018). Pleiotropic roles for ZEB1 in cancer. Cancer Res. 78, 30–35.

Chausovsky, A., Bershadsky, A.D., and Borisy, G.G. (2000). Cadherin-mediated regulation of microtubule dynamics. Nat. Cell Biol. 2, 797–804.

Chavez, K.J., Garimella, S. V., and Lipkowitz, S. (2010). Triple negative breast cancer cell lines: One tool in the search for better treatment of triple negative breast cancer. Breast Dis. 32, 35–48.

Combs, J., Kim, S.J., Tan, S., Ligon, L.A., Holzbaur, E.L., Kuhn, J., and Poenie, M. (2006). Recruitment of dynein to the Jurkat immunological synapse. Proc Natl Acad Sci U S A 103, 14883–14888.

Das, R.M., and Storey, K.G. (2014). Apical abscission alters cell polarity and dismantles the primary cilium during neurogenesis. Science (80-.). 343, 200–204.

Dent, R., Hanna, W.M., Trudeau, M., Rawlinson, E., Sun, P., and Narod, S.A. (2009). Pattern of metastatic spread in triple-negative breast cancer. Breast Cancer Res. Treat. 115, 423–428.

Dongre, A., and Weinberg, R.A. (2019). New insights into the mechanisms of epithelial–mesenchymal transition and implications for cancer. Nat. Rev. Mol. Cell Biol. 20, 69–84.

Doyle, A.D., Wang, F.W., Matsumoto, K., and Yamada, K.M. (2009). One-dimensional topography underlies three-dimensional fibrillar cell migration. J. Cell Biol. 184, 481–490.

Duchemin-Pelletier, E., Baulard, M., Spreux, E., Prioux, M., Burute, M., Mograbi, B., Guyon, L., Théry, M., Cochet, C., and Filhol, O. (2017). Stem cell-like properties of CK2β-down regulated mammary cells. Cancers (Basel). 9.

Duong, T., Goud, B., and Schauer, K. (2012). Closed-form density-based framework for automatic detection of cellular morphology changes. Proc. Natl. Acad. Sci. U. S. A. 2012.

Dupin, I., Camand, E., and Etienne-Manneville, S. (2009). Classical cadherins control nucleus and centrosome position and cell polarity. J. Cell Biol. 185, 779–786.

Elric, J., and Etienne-Manneville, S. (2014). Centrosome positioning in polarized cells: Common themes and variations. Exp. Cell Res. 1–9.

de Forges, H., Bouissou, A., and Perez, F. (2012). Interplay between microtubule dynamics and intracellular organization. Int. J. Biochem. Cell Biol. 44, 266–274.

Gauthaman, K., Fong, C.Y., and Bongso, A. (2010). Effect of ROCK inhibitor Y-27632 on normal and variant human embryonic stem cells (hESCs) in vitro: Its benefits in hESC expansion. Stem Cell Rev. Reports 6, 86–95.

Godinho, S.A., Picone, R., Burute, M., Dagher, R., Su, Y., Leung, C.T., Polyak, K., Brugge, J.S., Théry, M., and Pellman, D. (2014). Oncogene-like induction of cellular invasion from centrosome amplification. Nature 510.

Gomes, E.R., Jani, S., and Gundersen, G.G. (2005). Nuclear movement regulated by Cdc42, MRCK, myosin, and actin flow establishes MTOC polarization in migrating cells. Cell 121, 451–463.

Grigoriadis, A., Mackay, A., Noel, E., Wu, P., Natrajan, R., Frankum, J., Reis-Filho, J.S., and Tutt, A. (2012). Molecular characterisation of cell line models for triple-negative breast cancers. BMC Genomics 13, 619.

Gubelmann, C., Schwalie, P.C., Raghav, S.K., Röder, E., Delessa, T., Kiehlmann, E., Waszak, S.M., Corsinotti, A., Udin, G., Holcombe, W., et al. (2014). Identification of the transcription factor ZEB1 as a central component of the adipogenic gene regulatory network. Elife 3, 1–30.

Gupta, P.B., Pastushenko, I., Skibinski, A., Blanpain, C., and Kuperwasser, C. (2019). Phenotypic Plasticity: Driver of Cancer Initiation, Progression, and Therapy Resistance. Cell Stem Cell 24, 65–78.

Holenstein, C.N., Horvath, A., Schër, B., Schoenenberger, A.D., Bollhalder, M., Goedecke, N., Bartalena, G., Otto, O., Herbig, M., Guck, J., et al. (2019). The relationship between metastatic potential and in vitro mechanical properties of osteosarcoma cells. Mol. Biol. Cell 30, 887–898.

Huang, J., Huang, L., Chen, Y.-J., Austin, E., Devor, C.E., Roegiers, F., and Hong, Y. (2011). Differential regulation of adherens junction dynamics during apical-basal polarization. J. Cell Sci.

Hudis, C.A., and Gianni, L. (2011). Triple-Negative Breast Cancer: An Unmet Medical Need. Oncologist 16, 1–11.

Jang, M.H., Kim, H.J., Kim, E.J., Chung, Y.R., and Park, S.Y. (2015). Expression of epithelial-mesenchymal transition-related markers in triple-negative breast cancer: ZEB1 as a potential biomarker for poor clinical outcome. Hum. Pathol. 46, 1267–1274.

Jones, D.H., Gray, E.G., and Barron, J. (1980). Cold stable microtubules in brain studied in fractions and slices. J. Neurocytol. 9, 493–504.

Kardassis, D., Murphy, C., Fotsis, T., Moustakas, A., and Stournaras, C. (2009). Control of transforming growth factor signal transduction by small GTPases. FEBS J. 276, 2947–2965.

Kasioulis, I., Das, R.M., and Storey, K.G. (2017). Inter-dependent apical microtubule and actin dynamics orchestrate centrosome retention and neuronal delamination. Elife 6, e26215.

Kelava, I., and Lancaster, M.A. (2016). Stem Cell Models of Human Brain Development. Cell Stem Cell 18, 736–748.

Kraning-Rush, C.M., Califano, J.P., and Reinhart-King, C. a (2012). Cellular traction stresses increase with increasing metastatic potential. PLoS One 7.

Krebs, A.M., Mitschke, J., Losada, M.L., Schmalhofer, O., Boerries, M., Busch, H., Boettcher, M., Mougiakakos, Di., Reichardt, W., Bronsert, P., et al. (2017). The EMT-activator Zeb1 is a key factor for cell plasticity and promotes metastasis in pancreatic cancer. Nat. Cell Biol. 19, 518–529.

Kröger, C., Afeyan, A., Mraz, J., Eaton, E.N., Reinhardt, F., Khodor, Y.L., Thiru, P., Bierie, B., Ye, X., Burge, C.B., et al. (2019). Acquisition of a hybrid E/M state is essential for tumorigenicity of basal breast cancer cells. Proc. Natl. Acad. Sci. 201812876.

Laurent, J., Blin, G., Chatelain, F., Vanneaux, V., Fuchs, A., Larghero, J., and Théry, M. (2017). Convergence of microengineering and cellular self-organization towards functional tissue manufacturing. Nat. Biomed. Eng. 1.

Leal-Egaña, A., Letort, G., Martiel, J.-L., Christ, A., Vignaud, T., Roelants, C., Filhol, O., and Théry, M. (2017). The size-speed-force relationship governs migratory cell response to tumorigenic factors. Mol. Biol. Cell 28.

Lele, T.P., Dickinson, R.B., and Gundersen, G.G. (2018). Mechanical principles of nuclear shaping and positioning. J. Cell Biol. 217, 3330–3342.

Letort, G., Nedelec, F., Blanchoin, L., and Thery, M. (2016). Centrosome centering and decentering by microtubule network rearrangement. Mol. Biol. Cell 27, 2833–2843.

Libanje, F., Raingeaud, J., Luan, R., Thomas, Z.A., Zajac, O., Veiga, J., Marisa, L., Adam, J., Boige, V., Malka, D., et al. (2019). ROCK 2 inhibition triggers the collective invasion of colorectal adenocarcinomas. 1–23.

Ligon, L.A., Karki, S., Tokito, M., and Holzbaur, E.L.F. (2001). Dynein binds to β -catenin and may tether microtubules at adherens junctions. Nat. Cell Biol. 3, 913–917.

Lindley, L.E., and Briegel, K.J. (2010). Molecular characterization of TGFbeta-induced epithelial-mesenchymal transition in normal finite lifespan human mammary epithelial cells. Biochem. Biophys. Res. Commun. 399, 659–664.

Liu, Y., Lu, X., Huang, L., Wang, W., Jiang, G., Dean, K.C., Clem, B., Telang, S., Jenson, A.B., Cuatrecasas, M., et al. (2014). Different thresholds of ZEB1 are required for Ras-mediated tumour initiation and metastasis. Nat. Commun. 5, 1–9.

Luxton, G.W.G., Gomes, E.R., Folker, E.S., Vintinner, E., and Gundersen, G.G. (2010). Linear arrays of nuclear envelope proteins harness retrograde actin flow for nuclear movement. Science 329, 956–959.

Martiel, J.-L., Leal, A., Kurzawa, L., Balland, M., Wang, I., Vignaud, T., Tseng, Q., and Théry, M. (2015). Measurement of cell traction forces with ImageJ. Methods Cell Biol. 125, 269–287.

McBeath, R., Pirone, D.M., Nelson, C.M., Bhadriraju, K., and Chen, C.S. (2004). Cell shape, cytoskeletal tension, and RhoA regulate stem cell lineage commitment. Dev. Cell 6, 483–495.

Meng, W., Mushika, Y., Ichii, T., and Takeichi, M. (2008). Anchorage of microtubule minus ends to adherens junctions regulates epithelial cell-cell contacts. Cell 135, 948–959.

Mierke, C.T., Frey, B., Fellner, M., Herrmann, M., and Fabry, B. (2011). Integrin α5β1 facilitates cancer cell invasion through enhanced contractile forces. J. Cell Sci. 124, 369–383.

Mimori-Kiyosue, Y. (2011). Shaping microtubules into diverse patterns: Molecular connections for setting up both ends. Cytoskeleton 68, 603–618.

Morel, A.-P., Ginestier, C., Pommier, R.M., Cabaud, O., Ruiz, E., Wicinski, J., Devouassoux-Shisheboran, M., Combaret, V., Finetti, P., Chassot, C., et al. (2017). A stemness-related ZEB1–MSRB3 axis governs cellular pliancy and breast cancer genome stability. Nat. Med. 23, 568–578.

Morel, A.P., Hinkal, G.W., Thomas, C., Fauvet, F., Courtois-Cox, S., Wierinckx, A., Devouassoux-Shisheboran, M., Treilleux, I., Tissier, A., Gras, B., et al. (2012). EMT inducers catalyze malignant transformation of mammary epithelial cells and drive tumorigenesis towards claudin-low tumors in transgenic mice. PLoS Genet. 8.

Nieto, M.A., Huang, R.Y.-J., Jackson, R.A., and Thiery, J.P. (2016). Emt: 2016. Cell 166, 21–45.

Nigg, E.A. (2014). Centrosomes as signalling centres. Philos. Trans. R. Soc. Lond. B. Biol. Sci. 369.

Pakzad, M., Totonchi, M., Taei, A., Seifinejad, A., Hassani, S.N., and Baharvand, H. (2010). Presence of a ROCK inhibitor in extracellular matrix supports more undifferentiated growth of feeder-free human embryonic and induced pluripotent stem cells upon passaging. Stem Cell Rev. Reports 6, 96–107.

Pastushenko, I., and Blanpain, C. (2019). EMT Transition States during Tumor Progression and Metastasis. Trends Cell Biol. 29, 212–226.

Pastushenko, I., Brisebarre, A., Sifrim, A., Fioramonti, M., Revenco, T., Boumahdi, S., Van Keymeulen, A., Brown, D., Moers, V., Lemaire, S., et al. (2018). Identification of the tumour transition states occurring during EMT. Nature 556, 463–468.

Paszek, M.J., Zahir, N., Johnson, K.R., Lakins, J.N., Rozenberg, G.I., Gefen, A., Reinhart-King, C. a, Margulies, S.S., Dembo, M., Boettiger, D., et al. (2005). Tensional homeostasis and the malignant phenotype. Cancer Cell 8, 241–254.

Pitaval, A., Senger, F., Letort, G., Gidrol, X., Guyon, L., Sillibourne, J., and Théry, M. (2017). Microtubule stabilization drives 3D centrosome migration to initiate primary ciliogenesis. J. Cell Biol. 216.

Puisieux, A., Pommier, R.M., Morel, A., and Lavial, F. (2018). Cellular Pliancy and the Multistep Process of Tumorigenesis. Cancer Cell 33, 164–172.

Rodriguez-Fraticelli, A.E., Auzan, M., Alonso, M. a, Bornens, M., and Martin-Belmonte, F. (2012). Cell confinement controls centrosome positioning and lumen initiation during epithelial morphogenesis. J. Cell Biol. 198, 1011–1023.

Rodriguez-Hernandez, I., Cantelli, G., Bruce, F., and Sanz-Moreno, V. (2016). Rho, ROCK and actomyosin contractility in metastasis as drug targets. F1000Research 5, 783.

Schauer, K., Duong, T., Bleakley, K., Bardin, S., Bornens, M., and Goud, B. (2010). Probabilistic density maps to study global endomembrane organization. Nat. Methods 7, 560–566.

Schmoranzer, J., Fawcett, J.P., Segura, M., Tan, S., Vallee, R.B., Pawson, T., and Gundersen, G.G. (2009). Par3 and dynein associate to regulate local microtubule dynamics and centrosome orientation during migration. Curr. Biol. 19, 1065–1074.

Shahbazi, M.N., Megias, D., Epifano, C., Akhmanova, A., Gundersen, G.G., Fuchs, E., and Perez-Moreno, M. (2013). CLASP2 interacts with p120-catenin and governs microtubule dynamics at adherens junctions. J. Cell Biol. 203, 1043–1061.

Sipe, C.W., Liu, L., Lee, J., Grimsley-Myers, C., and Lu, X. (2013). Lis1 mediates planar polarity of auditory hair cells through regulation of microtubule organization. Development 140, 1785–1795.

Spaderna, S., Schmalhofer, O., Wahlbuhl, M., Dimmler, A., Bauer, K., Sultan, A., Hlubek, F., Jung, A., Strand, D., Eger, A., et al. (2008). The Transcriptional Repressor ZEB1 Promotes Metastasis and Loss of Cell Polarity in Cancer. Cancer Res. 68, 537–544.

Stehbens, S.J., Paterson, A.D., Crampton, M.S., Shewan, A.M., Ferguson, C., Akhmanova, A., Parton, R.G., and Yap, A.S. (2006). Dynamic microtubules regulate the local concentration of E-cadherin at cell-cell contacts. J Cell Sci 119, 1801–1811.

Stemmler, M.P., Eccles, R.L., Brabletz, S., and Brabletz, T. (2019). Non-redundant functions of EMT transcription factors. Nat. Cell Biol. 21, 102–112.

Stinchcombe, J.C., and Griffiths, G.M. (2014). Communication, the centrosome and the immunological synapse. Philos. Trans. R. Soc. Lond. B. Biol. Sci. 369, 20130463-.

Tang, N., and Marshall, W.F. (2012). Centrosome positioning in vertebrate development. J. Cell Sci. 125, 4951–4961.

Tavares, A.L.P., Mercado-Pimentel, M.E., Runyan, R.B., and Kitten, G.T. (2006). TGFβ-mediated RhoA expression is necessary for epithelial-mesenchymal transition in the embryonic chick heart. Dev. Dyn. 235, 1589–1598.

Tavares, S., Vieira, A.F., Taubenberger, A.V., Araújo, M., Martins, N.P., Brás-Pereira, C., Polónia, A., Herbig, M., Barreto, C., Otto, O., et al. (2017). Actin stress fiber organization promotes cell stiffening and proliferation of pre-invasive breast cancer cells. Nat. Commun. 8.

Théry, M. (2010). Micropatterning as a tool to decipher cell morphogenesis and functions. J. Cell Sci. 123, 4201–4213.

Théry, M., Racine, V., Piel, M., Pépin, A., Dimitrov, A., Chen, Y., Sibarita, J.-B., and Bornens, M. (2006). Anisotropy of cell adhesive microenvironment governs cell internal organization and orientation of polarity. Proc. Natl. Acad. Sci. U. S. A. 103, 19771–19776.

Thiery, J.P., Acloque, H., Huang, R.Y.J., and Nieto, M.A. (2009). Epithelial-mesenchymal transitions in development and disease. Cell 139, 871–890.

Tseng, Q., Wang, I., Duchemin-Pelletier, E., Azioune, A., Carpi, N., Gao, J., Filhol, O., Piel, M., Théry, M., and Balland, M. (2011). A new micropatterning method of soft substrates reveals that different tumorigenic signals can promote or reduce cell contraction levels. Lab Chip 11, 2231–2240.

Tseng, Q., Duchemin-Pelletier, E., Deshiere, A., Balland, M., Guillou, H., Filhol, O., and Théry, M. (2012). Spatial organization of the extracellular matrix regulates cell-cell junction positioning. Proc. Natl. Acad. Sci. U. S. A. 109, 1506–1511.

Tsubakihara, Y., and Moustakas, A. (2018). Epithelial-Mesenchymal Transition and Metastasis under the Control of Transforming Growth Factor β. Int. J. Mol. Sci. 19, 1–30.

Uhler, C., and Shivashankar, G. V. (2017). Regulation of genome organization and gene expression by nuclear mechanotransduction. Nat. Rev. Mol. Cell Biol.

Vasileva, E., and Citi, S. (2018). The role of microtubules in the regulation of epithelial junctions. Tissue Barriers 6, 1–20.

Vianello, S., and Lutolf, M.P. (2019). Understanding the Mechanobiology of Early Mammalian Development through Bioengineered Models. Dev. Cell 48, 751–763.

Vignaud, T., Ennomani, H., and Théry, M. (2014). Polyacrylamide Hydrogel Micropatterning.

Wan, A.C.A. (2016). Recapitulating Cell–Cell Interactions for Organoid Construction – Are Biomaterials Dispensable? Trends Biotechnol. xx, 1–11.

Yoshida, T., Ozawa, Y., Kimura, T., Sato, Y., Kuznetsov, G., Xu, S., Uesugi, M., Agoulnik, S., Taylor, N., Funahashi, Y., et al. (2014). Eribulin mesilate suppresses experimental metastasis of breast cancer cells by reversing phenotype from epithelial–mesenchymal transition (EMT) to mesenchymal– epithelial transition (MET) states. Br. J. Cancer 110, 1497–1505.

Yu, M., Bardia, A., Wittner, B.S., Stott, S.L., Smas, M.E., Ting, D.T., Isakoff, S.J., Ciciliano, J.C., Wells, M.N., Shah, A.M., et al. (2013). Circulating breast tumor cells exhibit dynamic changes in epithelial and mesenchymal composition. Science 339, 580–584.

